# Exploring mechanisms of scar-free skin wound healing in adult zebrafish in comparison to mouse

**DOI:** 10.1101/2025.09.09.675222

**Authors:** İsmail Küçükaylak, Francisco Javier Martínez Morcillo, Kai Halwas, Nils Reiche, Manuel Metzger, Petra Comelli, Jürgen Brinckmann, Sabine Eming, Matthias Hammerschmidt

**Author notes:** Department of Biology, Institute of Molecular Health Sciences, ETH Zurich, Zurich, Switzerland. Departamento de Biología Celular e Histología, Facultad de Biología, Universidad de Murcia, Murcia, Spain (Francisco Javier Martínez Morcillo).

## Abstract

Adult zebrafish have the ability to perfectly regenerate their skin after injury without leaving a scar behind. Yet, they intermediately form a collagen-rich granulation tissue that later fully regresses. In contrast, adult mammals lose this ability, resulting in persistent tissue fibrosis and scarring. We performed single-cell RNA sequencing to better characterize the dynamics and heterogeneity of involved cell types during different stages of zebrafish cutaneous wound healing, focusing on macrophages and fibroblasts. Macrophage subclusters display pro- and/or anti-inflammatory/repair characteristics, and fibroblast subclusters characteristics of extracellular matrix formation and degradation, which largely co-exist during all stages of wound healing. Strikingly, some fibroblasts display signatures of myofibroblasts, implicated in fibrotic healing in mammals. In addition, zebrafish fibroblasts express multiple genes with described pro-fibrotic effects in mammalian models. One of them is *plod2*, which encodes lysylhydroxylase 2. In cutaneous mouse wounds, *Plod2* is induced in fibroblasts by macrophage-released Resistin-like molecule RELMα encoded by the *Retlna* gene, promoting the formation of DHLNL collagen crosslinks and thereby less resolvable fibrotic tissue. *retln* genes are absent from the zebrafish genome; nevertheless, *plod2* expression is initiated in zebrafish dermal fibroblasts upon wounding, in this case via TGFβ signaling, accompanied by increased collagen DHLNL crosslinking. Yet, both transgenic overexpression and genetic knock-out of *plod2* do not interfere with granulation tissue formation and regression, pointing to additional pathways assuring the resolution of transient fibrosis in zebrafish skin wounds even in the presence of strong collagen crosslinking.

## Introduction

Regeneration is defined by the almost perfect restoration of the lost or injured structure while repair is defined by an inflammatory response, connective tissue and finally scar formation, which is also observed in adult mammalian skin wound healing [1]. The healing of wounded skin upon injury is critical and skin integrity should be restored again swiftly since it constitutes the barrier to the outside world. Interestingly, adult zebrafish are able to fully regenerate their skin upon external insults, whereas mammalian scar-free regeneration is lost during embryogenesis [2–4]. Therefore, it is important to understand the mechanisms underlying this process in order to treat wound healing-related diseases properly.

The injury response initiates the formation of a new stroma underneath the neo-epidermis called the granulation tissue, where fibroblasts, innate immune cells and blood vessels invade the wounded area [5]. Macrophages, one of the main players during cutaneous wound healing, have essential roles by determining regeneration or scar tissue formation. Shortly after injury, tissue-resident macrophages are complemented by inflammatory monocytes that enter the damaged tissue to give rise to macrophages which clear cell debris and damaged extracellular matrix (ECM), while attracting other immune cells through the secretion of pro-inflammatory cytokines such as tumor necrosis factor α (TNF-α) and interleukin-1 (IL-1β). Later, they also play a critical role in resolving inflammation and to promote ECM tissue reconstruction, cell proliferation, and angiogenesis. These later macrophages are characterized by the secretion of anti-inflammatory factors such as IL-10 and factors like transforming growth factor β (TGF-β) that promote tissue repair [6–8]. These early and late functions are most likely carried out by distinct macrophage subpopulations [9, 10], with an underlying macrophage plasticity and heterogeneity that turned out as much more complex than the initial concept of early pro-inflammatory, classically activated M1 macrophages and late alternatively activated M2 macrophages reducing inflammation and promoting tissue repair [11–13].

Similar to macrophages, fibroblasts of the granulation tissue of mammalian wounds have been reported to stem from different sources [14–17]. Fibroblasts residing in the adjacent dermis proliferate and migrate into the wound bed. Furthermore, monocytes entering the wound from the blood can differentiate into fibroblast-like cells called fibrocytes [18] – as also present in lesions associated with multiple fibrosing diseases [19, 20]. Within the granulation tissue, dermal fibroblasts are stimulated by growth factors like TGFβ-1 and mechanical stress to synthesize and secrete collagens, which are the main constituents of the wound ECM, as well as other ECM molecules such as fibronectin, proteoglycans, glycosaminoglycans and hyaluronic acid [21]. Some fibroblasts are further stimulated to differentiate into myofibroblasts that in addition to ECM production form intracellular stress fibers and muscle proteins in order to establish tissue contraction [22]. The differentiation of fibroblasts to myofibroblasts during mammalian wound healing has also been described as a key feature for fibrosis; while they are beneficial for physiological tissue remodelling, they have adverse effects when becoming excessive, such as during hypertrophic scarring of wounded tissue [22, 23]. However, not all, but particular fibroblast lineages / subclusters of the granulation tissue are supposed to develop such adverse features [17, 24–28], dependent on corresponding inflammatory priming which dictates on regenerative versus fibrotic repair [15, 29].

The general consequence of fibrosis is an aberrant accumulation of ECM components and the loss of a normal tissue architecture [30–32]. In mammalian wounds, fibrosis denotes persistent excessive scar tissue, whereas in zebrafish wounds, cutaneous fibrosis is only transient and the original tissue structure is fully restored. An important factor contributing to collagen stability/persistence within the granulation tissue is the mode of collagen crosslinking via lysine residues that are already accordingly modified within the ER of fibroblasts. Lysylhydroxylase 2 (LH2) encoded by the *PLOD2* gene specifically hydroxylates lysine residues in the telopeptides of collagen polypeptides, which is an essential step for the subsequent Lysyloxidase (LOX)-mediated formation of dihydroxy lysinonorleucine (DHLNL) cross-links of collagen fibrils [33–36]. Such DHLNL cross-links are supposed to make collagens less susceptible to degradation and are a typical feature of mechanically stiff tissues such as bone and cartilage, whereas soft connective tissues such as normal skin are characterized by LH2-independent, lysinealdehyde-derived collagen cross-links (hydroxylysinonorleucine, HLNL) [37, 38]. Mouse skin wounds, on the other side, have been reported to display increased LH2 levels as well as increased DHLNL collagen cross-linking, while an alleviation of wound LH2 levels by genetic ablation of RELMα, a fibroblast-stimulating factor made by wound macrophages, reduced DHLNL levels as well as scarring [39]. Together, this points to an essential pro-fibrotic effect of LH2 during the scarring of mammalian skin wounds.

Here, we investigated the cellular and the molecular dynamics involved in zebrafish scar-free cutaneous wound healing. Utilizing scRNA sequencing, we analyzed unwounded skin and various stages of wound healing, with particular focus on the roles and interactions of macrophages and fibroblasts. Our findings highlight the heterogeneity of these cell populations, with zebrafish dermal fibroblast subclusters actually expressing multiple genes associated with fibrosis in mammalian systems, including *plod2*, but also genes with no annotated function yet. Interestingly, neither gain nor loss of function of *plod2* altered the regenerative ability of zebrafish, suggesting that alternative factors are involved that make the difference to scarring in mammals. Collectively, our data set the base for systematic comparative analysis between perfectly regenerating and imperfectly healing model organisms as a promising approach to identify new factors that determine the quality of the reparative outcome.

## Results

### scRNA-seq-based identification and quantification of different cell types during different stages of cutaneous wound healing in adult zebrafish

We have formerly shown that full-thickness wounds in the skin of adult zebrafish introduced by a dermatology laser removing the epidermis, dermis, scales and subcutaneous adipocytes, fully close again within 10 hours after wounding. This closure is purely driven by morphogenetic movements within the epidermis, with keratinocytes starting to re-epithelialize the injured area even before innate immune cells have entered the wound bed [4, 40]. However, inflammation by neutrophils and macrophages is required for the later recruitment of fibroblasts and the formation of a collagen-rich granulation tissue underneath the neo-epidermis [4]. We further showed that in untreated fish, wound inflammation is maximal at two days post wounding (dpw), when granulation tissue starts to form (Fig 1B), that granulation tissue is maximal at 4 dpw (Fig 1C), and that granulation tissue is almost completely resolved at 8 dpw, followed by the regeneration skin appendages like scales [4], in sum constituting perfect and scar-free skin healing with only temporary fibrosis.

**Fig 1.**
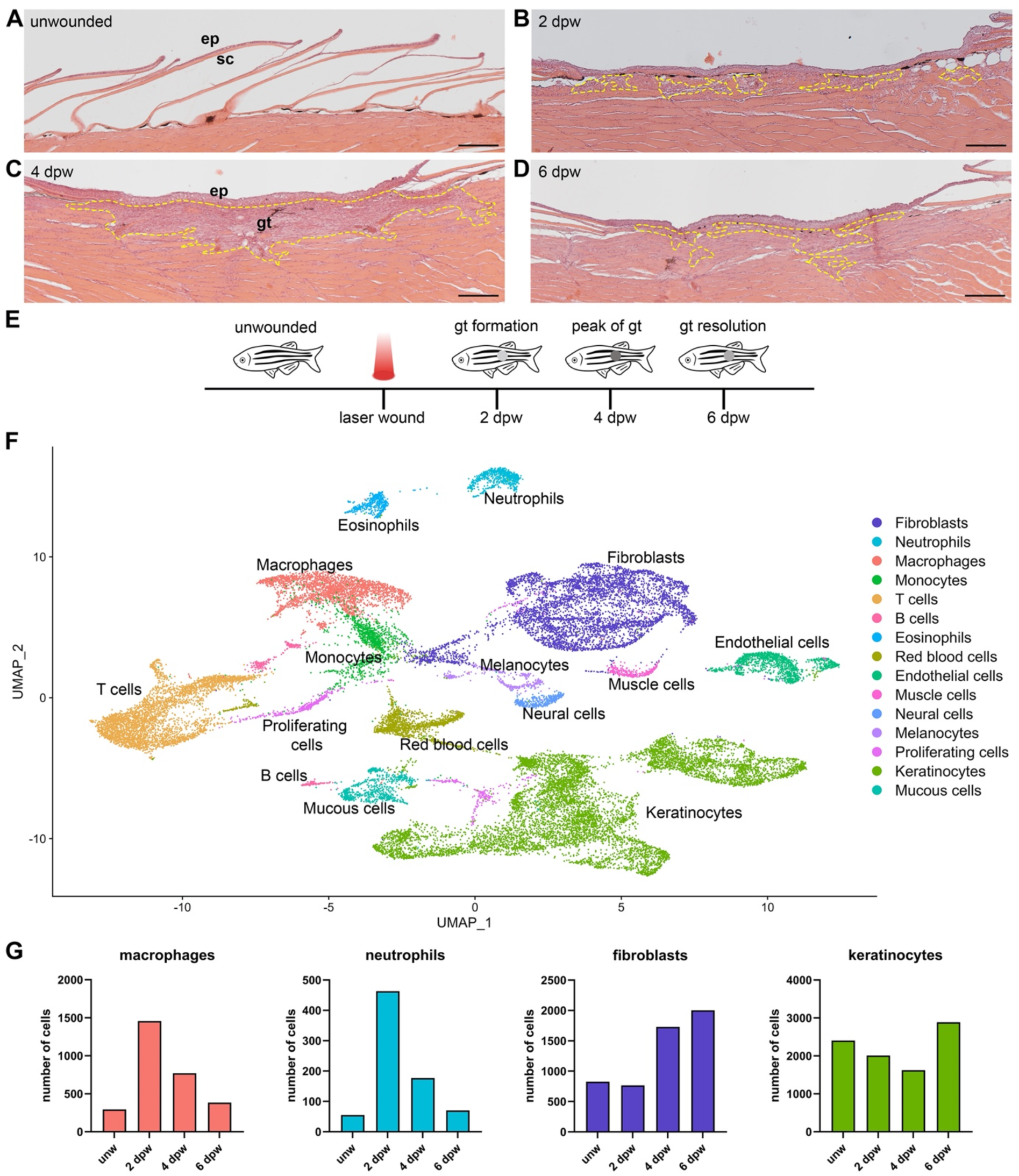
Single cell transcriptome analysis reveals the cellular composition of zebrafish cutaneous wounds. (A-D) Hematoxylin and eosin (H&E) staining of longitudinal skin sections showing the formation of granulation tissue (gt) beneath the wound, reaches the maximum size in volume at 4 days post wounding (dpw) and then regresses (yellow dashed lines mark the granulation tissue). (E) Cartoon for the experimental design. Unwounded skin tissue and biopsies of wounds from 2, 4 and 6 dpw were collected for single cell RNA sequencing. (F) UMAP representation of single cell RNA sequencing data after integration of four datasets and cell clustering results. (G) Absolute number of macrophages, neutrophils, fibroblasts and keratinocytes across the time points according to single cell RNA sequencing data set. Numerical values for panel G can be found in S1 Data. Scale bars: A-D = 200 µm. ep: epidermis.

To better understand the cellular dynamics underlying this reversible fibrosis, we performed comparative single-cell RNA-sequencing (scRNA-seq) of skin biopsies from unwounded skin (Fig 1A) and from skin wounds at 2 dpw, when granulation tissue has started to form (Fig 1B), at 4 dpw, when granulation tissue size peaks (Fig 1C) and at 6 dpw, when granulation tissue regresses (Fig 1D,E). scRNA-seq analysis and Uniform Manifold Approximation and Projection (UMAP) representation of all time points integrated revealed the different major cell types present during zebrafish cutaneous wound healing (Fig 1F; S1 Fig.). The top 10 most highly genes in the specific clusters were used to identify the cell types at the four investigated time points (S2 Fig.). The majority of the wound cells were comprised of keratinocytes, fibroblasts and immune cells. In line with former histological data [4], scRNA-seq revealed a significant increase in numbers of wound macrophages (5.6 fold) and neutrophils (8.4 fold) at 2 dpw compared to unwounded skin, with numbers progressively dropping again at 4 dpw and 6 dpw, although remaining higher than in unwounded skin (Fig 1G). In contrast, fibroblasts only displayed increased numbers compared to unwounded skin at 4 dpw, and even remained high at 6 dpw (Fig 1G). The latter might indicate that the reduction of granulation tissue from 4 to 6 dpw (Fig 1C,D) is mainly due to the degradation of ECM, rather than the loss of fibroblasts. In addition, it is possible that fibroblasts are, in fact, lost – according to our former *col1a2* in situ hybridizations [4] and TUNEL-stainings [41] most likely via fibroblast emigration, rather than cell death - and that such wound-adjacent emigrating fibroblasts were still contained in our scRNA-seq samples.

### Cellular heterogeneity of wound macrophages

Data obtained for cutaneous wounds in mouse [9, 10, 24, 42–44] as well as wounds of other tissues in zebrafish [45–49] point to a large degree of cellular heterogeneity among both macrophages and fibroblasts. We therefore next analyzed macrophage and fibroblast clusters of our zebrafish skin wounds in more detail. For macrophages, we identified seven different subclusters (Fig 2A; S3 Fig.). Subclusters 0-5 were present at all four investigated stages, including unwounded skin. In contrast, subcluster 6 was only found at 2 dpw and 4 dpw, but was absent from unwounded skin and at 6 dpw, pointing to a specific role during (temporary) formation of new granulation tissue, rather than regular dermal homeostasis and granulation tissue regression (Fig 2B).

**Fig 2.**
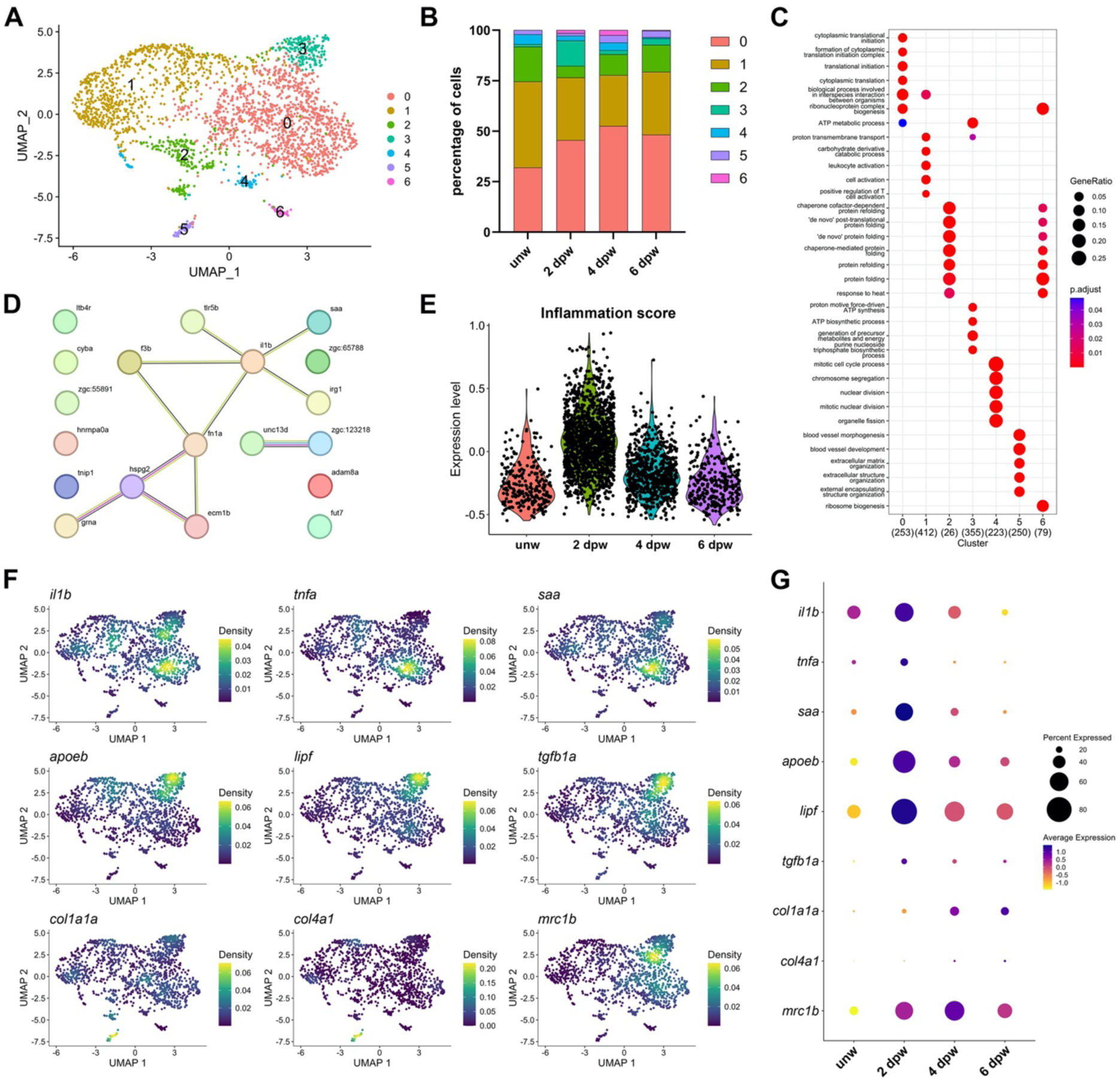
Characterization of distinct macrophage subpopulations. (A) UMAP representation of macrophage subclusters of four datasets. (B) Percentage of macrophage subcluster numbers across the time points. Numerical values for panel B can be found in S1 Data. (C) Gene ontology (GO) analysis of macrophage subclusters at 2 days post wounding (dpw). The GeneRatios, indicated by the diameter of dots, represent the proportion of input genes associated with a given GO term, calculated as the number of genes annotated to the term divided by the total number of input genes used in the analysis; colors represent the significance of the enrichment (p.adjust). (D) STRING analysis [128] of genes up-regulated in macrophages at 2 dpw in comparison with unwounded samples, yielding the shown pathway “GO:0006954-Inflammatory response”. (E) Violin plot of inflammation scores of macrophages in unwounded skin and at 2 dpw, 4 dpw and 6 dpw, calculated via module scoring using Seurat’s AddModuleScore function with the genes shown in panel D. (F) UMAP and density plot of the expression of some selected genes at 2 dpw for pro- and anti-inflammatory phenotypes of macrophages. (G) Dot plot showing the expression level of the selected genes across the time points. Circle size represents the percentage of cells expressing the genes and the colour represents the level of gene expression.

The different subclusters were further characterized via gene ontology (GO) enrichment analyses (Fig 2C), based on significantly upregulated genes (S4 Fig.), and by comparing the expression levels of genes (even if they were not among the top 10) with formerly reported pro-inflammatory, anti-inflammatory and/or tissue repair functions.

At 2 dpw, *il1b*, *tnfa* and *saa* genes, encoding interleukin 1, tumor necrosis factor α, and serum amyloid A, respectively, and associated with pro-inflammatory roles [50–54] were mainly expressed in cluster 0, whereas *apoeb*, *lipf*, *tgfb1a* genes, encoding apolipoprotein E, lipase F, and transforming growth factor β1, respectively, and associated with anti-inflammatory and repair/fibrosis-stimulating roles [10, 45, 51, 54], were mainly expressed in cluster 3 (Fig 2F). Of note, the expression of both *tnfa* in cluster 0 and *tgfb1a in cluster 3* dropped between 2 dpw and 4 dpw (S5 Fig.). For TNFα, this is in line with its potential early proinflammatory role, while for TGFβ, its potential later anti-inflammatory/pro-repair task might be taken over by the concurrent up-regulation of *tgfb1a* expression in fibroblasts (see below). Expression of *il10*, a strong anti-inflammatory marker and regulator in mammalian wounds, was only detected in unwounded skin, but not in macrophages of any subcluster and at any stage of our zebrafish wounds. Pro-inflammatory cluster 0 was also present at 4 dpw and 6 dpw (Fig 2B,G), although the overall inflammation score was maximal at 2 dpw and dropped back to unwounded skin levels during later stages (Fig 2E; see legends of panels D and E for more details). Interestingly, cluster 0, in addition to its pro-inflammatory role might also play an anti-fibrotic, ECM-degrading role, as suggested by the strong and continuous expression of *mrc1b* (Fig 2G, S3 Fig., S5 Fig.), encoding a mannose receptor formerly shown to be required for collagen phagocytosis and degradation by M2-like macrophages in the mouse skin [55].

Cluster 1, the only cluster with a high score for GO terms “leukocyte activation” and “cell activation” (Fig 2C), displayed strong expression of *cxcl19* (S3 Fig., S5 Fig.). It encodes a chemoattractant which in other physiological contexts has been reported to signal via CXCR4 receptors [56], while CXCR4 has been shown to be the predominant chemokine receptor on fibrocytes and to be involved in multiple pathological mechanisms in fibrosis [57]. Indeed, we found the two zebrafish homologs, *cxcr4a* and *cxcr4b*, also sparsely expressed in wound fibroblasts (S5 Fig.), pointing to a potential role of cluster 1 macrophages to attract fibroblasts to the wound. However, in particular *cxcr4b* was much more strongly expressed in macrophages themselves and in all other immune cell types (S5 Fig.), pointing to an additional global pro-inflammatory role of cluster 1.

Cluster 5 was particularly characterized by its high score for GO term “extracellular structure organization” (Fig 2C, S4 Fig.). Cluster 5 macrophages expressed for instance *col1a1a* (Fig 2F,G, S5 Fig.) encoding fibrillar collagen type I, which is usually and mainly produced by fibroblasts [4, 15, 58] (see below), as well as *col4a1, col4a2* and *lama4* (S3 Fig., S5 Fig.), encoding integral components of the skin basement membrane - which also has to regenerate at the epidermal-dermal junction, a function mainly performed by keratinocytes and fibroblasts. Interestingly, laminins and collagen IV have also been implicated with fibroblast activation and fibrosis in different mammalian systems [59]. Furthermore, macrophages have been formerly reported to directly contribute to collagen and ECM production upon heart injury in zebrafish and mouse [60]. Together, this suggests that macrophage subcluster 5 might, in addition to stimulating fibroblasts, directly contribute to ECM production during granulation tissue formation in zebrafish skin wounds.

Cluster 6, the only wound-specific of the seven macrophage subclusters (Fig 2B), was characterized by high expression levels of genes like *f13a1a*, *esr2a* and *cpn1* (S3 Fig., S5 Fig.). *f13a1a* encodes transglutaminase factor XIII-A, which is mainly involved in blood clotting – in contrast to mammals, fish skin wounds lack blood clots [4] – but which in mammals has also been described at the surface of alternatively activated M2 macrophages to terminate inflammation and to promote wound healing and phagocytosis [61]. *esr2a* encodes an estrogen receptor 2/beta. In mice, Esr2 has been described to be required in keratinocytes [62] and Esr1/alpha in macrophages [63] to allow proper cutaneous wound healing, while in zebrafish, it has been shown to accelerate heart regeneration [64]. Finally, *cpn1* encodes carboxypeptidase N, which to our knowledge had not been reported to be made by macrophages thus far, which, however, is known to inactivate bradykinin and anaphylatoxins such as C3a and C5a, thereby eliciting anti-inflammatory effects [65]. Furthermore, GO analysis demonstrated an enrichment of cluster 6 for de novo protein folding and distinctively for ribosome biogenesis (Fig 2C), pointing to strong protein-synthesizing activity. In sum, this points to cluster 6 as a wound healing-specific macrophage subpopulation with particular activation modes and a unique combination of anti-inflammatory and pro-regenerative effects.

Together, our macrophage subcluster analyses indicate that, in line with revised concepts in mammals [12], macrophages of zebrafish skin wounds do not follow the concept of stage-specific pro- and anti-inflammatory polarization, but that the different (and to some extent opposing) tasks fulfilled by macrophages are distributed among a whole spectrum of subpopulations that largely co-exist during the different stages of wound healing.

### Cellular heterogeneity of wound fibroblasts

Analysis of fibroblasts during the different stages of cutaneous wound healing revealed at total of six different subclusters (Fig 3A, S6 Fig.). Of all subclusters, cluster 0 showed the highest increase in cell numbers induced by wounding (from 8% of all fibroblasts in unwounded skin to 45% at 4 dpw; Fig 3B). In addition, one new wound-specific fibroblast subcluster not present in uninjured skin showed up, cluster 4, comprising 11% of all fibroblasts at 4 dpw (Fig 3B). Interestingly, parts of clusters 2 and 4 displayed a transcriptional signature pointing to myofibroblast-like cell functions, with specific expression of genes encoding muscle actin (*acta1a1*; Fig 3F,G, S8 Fig.) a marker of myofibroblasts in burned mammalian wounds [66], the muscle-specific intermediate filament desmin (*desma*; S8 Fig.), the muscle/myofibroblast-specific cell-adhesion molecule M-cadherin [67] (*cdh15*; S8 Fig.), or the Ankyrin repeat domain-containing protein 1 (*ankrd1a*; Fig 3F,G, S8 Fig.), also known as Cardiac ankyrin repeat protein (CARP), a stress-inducible myofibrillar protein and known target of TGFβ signaling, which, although mainly implicated in heart pathologies and carcinogenesis, was also reported to be elevated during hypertrophic skin scarring [68], while gene knockout led to reduced granulation tissue formation during cutaneous wound healing [69]. In mammalian skin wounds, myofibroblasts are considered as a crucial executer of fibrosis and scarring [23]. Of note, however, in zebrafish, we found these potential myofibroblasts, as well as entire fibroblast subcluster 4, not only to be absent from uninjured skin, but also – in contrast to all other fibroblast subclusters – to display a several-fold decrease of relative cell numbers during granulation tissue regression (6 dpw) compared to stages when granulation tissue is maximal (4 dpw) (Fig 3B, S8 Fig.).

**Fig 3.**
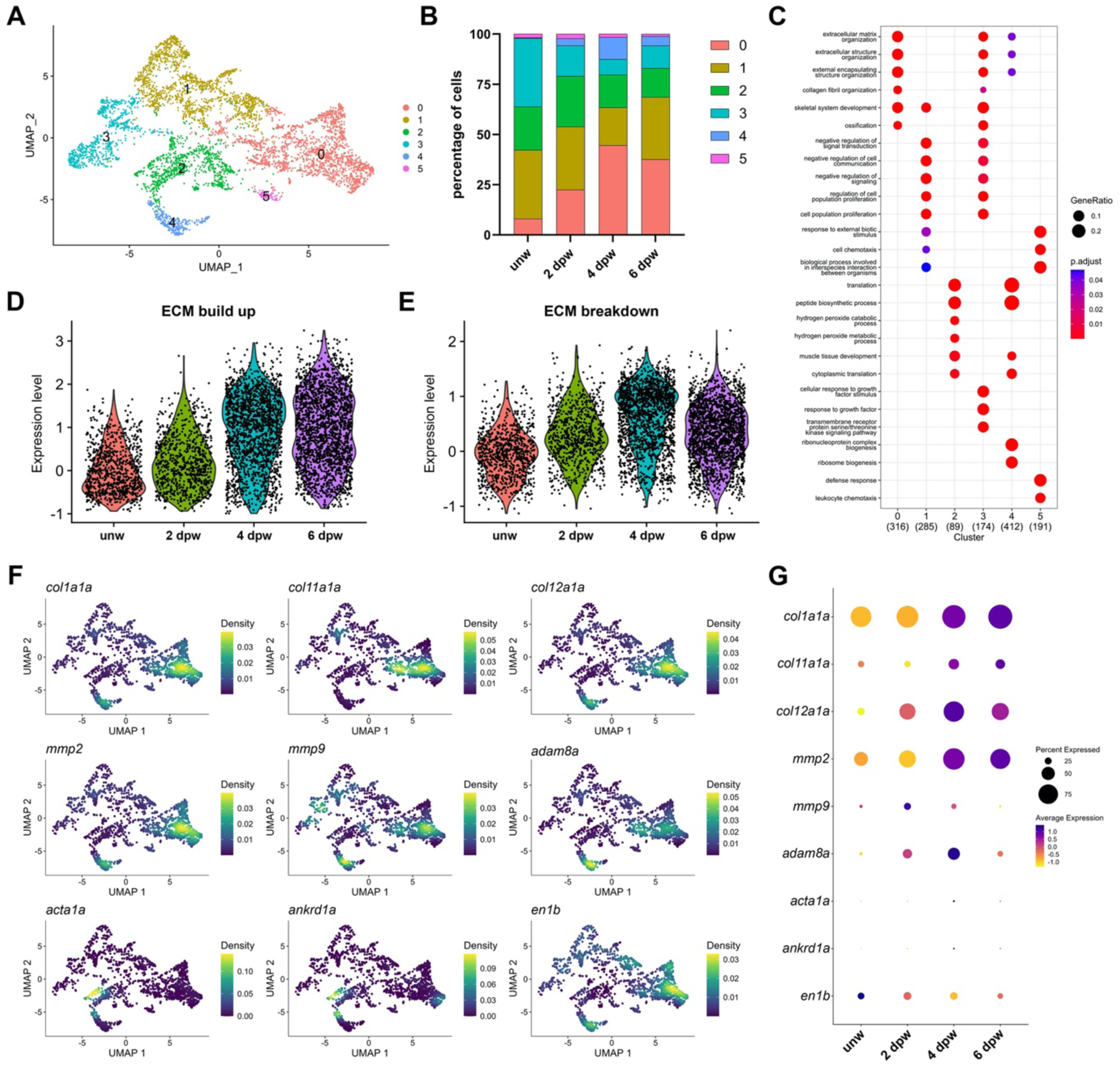
Characterization of distinct fibroblast subpopulations. (A) UMAP representation of fibroblast subclusters of four datasets. (B) Percentage of fibroblast subcluster numbers across the time points. Numerical values for panel B can be found in S1 Data. (C) Gene ontology (GO) analysis of fibroblast subclusters at 4 days post wounding (dpw). (D, E) Violin plots showing the expression level of ECM build up and ECM break down related genes in fibroblasts across the time points. (F) UMAP and density plot of the expression of some selected genes at 4 dpw, associated with ECM regulation or fibrosis. (G) Dot plot showing the expression level of the selected genes across the time points. Circle size represents the percentage of cells expressing the genes and the colour represents the level of gene expression.

Fibroblasts of zebrafish wounds also displayed expression of other genes marking fibroblast cell types implicated with mammalian fibrosis, for instance the Engrailed transcription factor gene *En1*. In mouse, En1 has been identified as a crucial pro-fibrotic factor that is highly up-regulated in a subset of wound fibroblasts upon skin injury, while ablation of these fibroblasts or blockage of En1 activation in them alleviated ECM production and prevented scarring [27, 28]. In fish skin, the En1 ortholog *en1b* was expressed rather broadly throughout the entire fibroblast cluster, while expression levels remained rather unaltered upon wounding and during the different phases of wound healing (S8 Fig.). A similarly broad expression within the entire fibroblast cluster, which was even strongly up-regulated upon wounding, was also observed for the two paralogs of mammalian Periostin (*postna*, *postnb*; S8 Fig.), a secreted protein that has for instance been shown to be strongly up-regulated in myofibroblasts of the injured mouse heart, while deletion of periostin-positive myofibroblasts reduced collagen production and scar formation [70].

GO enrichment analysis of the different zebrafish fibroblast subclusters at 4 dpw revealed “ECM organization” as a primary cellular function of clusters 0 and 3 and the wound-specific cluster 4 (Fig 3C, S6 Fig.). Of note, the three clusters differed in the collagen types [71] they primarily generate. *col1a1* and c*ol1a2*, encoding collagen I, the main dermal fibril-forming collagen, was produced by all three clusters (Fig 3C, S9 Fig.), while *col5a2b*, encoding collagen V, another fibrillar collagen that serves as a nucleator of collagen I fibril formation, was primarily produced by cluster 3 (S7 Fig., S9 Fig.), and *col12a1b*, encoding ColXII, a FACIT associated to collagen I-containing fibrils that has been implicated with pro-regenerative functions during zebrafish heart and spinal cord regeneration [47, 72], by clusters 0 and 4 (Fig 3C, S7 Fig., S9 Fig.). Interestingly, cluster 0 also generated fibrillar collagen II (*col2a1*) and fibrillar collagen XI (*col11a1b*) that serves as a nucleator of collagen II fibril formation (Fig 3C, S7 Fig., S9 Fig.). Such collagen II fibrils are mainly present in cartilage and have, at least to our knowledge, not been described in the context of mammalian cutaneous wound healing [24]; yet, administration of collagen type II isolated from sturgeon cartilage has been reported to improve the healing of cutaneous mouse wounds [73]. Cluster 3 was the only cluster that also displayed strong expression of *col10a1* (S7 Fig., S9 Fig.), encoding a network-forming collagen X that is mainly involved in hypertrophic cartilage and bone formation, osteoarthritis and, together with type I and type II collagens, in bone fracture healing [74–76], but has not been described in the context of skin wound healing. Yet, collagen X has been reported to serve as a ligand of the discoidin domain receptor DDR2 [77], and DDR2 has been shown to promote ECM remodeling and dermal wound healing in the mouse skin [78]. However, according to our own data, its expression is downregulated during stages of granulation tissue growth (2 dpw, 4 dpw; S9 Fig.), pointing to a role during dermal homeostasis and/or regression, rather than growth.

Cluster 4 was the only cluster also expressing *col18a1b* (S9 Fig.). It encodes collagen XVIII, a basement membrane proteoglycan that upon proteolytic cleavage for instance by matrix metalloproteases (MMPs) can give rise to endostatin, a matricryptin with multiple activities, including the blockage of MMP activation [79]. Mainly described in the context of angiogenesis and tumorigenesis, collagen XVIII and Endostatin have also been shown to have a negative impact on skin wound healing in mice [80], again in seeming contradiction to the increase of *col18a1b* expression levels during skin wound healing in zebrafish (S9 Fig.). Finally, *col4a1/2*, encoding network-forming basement membrane collagen IV, was primarily expressed in clusters 0 and 4 (S9 Fig.).

Similar to collagens, other ECM proteins were also generated by fibroblasts in a rather subcluster-specific manner. Genes encoding fibronectins (*fn1a/b*), which also form fibrils and like collagens can be cleaved by MMPs [81], were primarily expressed in clusters 0 and 4 (S10 Fig.). In addition, clusters 0 and 4 displayed strong expression of Cartilage Intermediate Layer Protein 1 (*cilp1*; S10 Fig.), a secreted ECM protein normally associated with bone and cartilage development, which also has been shown to bind TGFβ1, thereby suppressing TGFβ1-induced differentiation of fibroblasts to myofibroblasts during heart fibrosis [82]. Furthermore, cluster 0 was the only cluster displaying expression of osteoglycin (*ogna*; S10 Fig.), a small leucine-rich proteoglycan (SLRP) that has been shown to be negatively regulated by TGFβ and to block myofibroblast differentiation and fibrosis in different mammalian tissues, including the heart [83], thus, like *cilp1*, displaying clear anti-fibrotic effects. Similarly, wound-specific cluster 4 specifically expressed plasminogen activator, urokinase (uPA) (*plaub*; S10 Fig.), a secreted serine protease which, although widely upregulated in fibrotic diseases, has been shown to alleviate skin fibrosis [84].

ECM-degrading proteases displayed both subcluster-specific as well as more global fibroblast expression patterns. For example, the respectively secreted and membrane-tagged gelatinases *mmp2* and *mmp14b* [85], were broadly expressed in all fibroblast clusters, with higher levels at stages of granulation tissue regression (6 dpw) than at stages of granulation tissue formation, while the membrane-tagged disintegrin and metalloprotease 8 encoded by *adam8a* [86] was primarily expressed in clusters 0 and 4 (Fig 3F, S10 Fig.).

Of the fibroblast subclusters with primary cellular functions other than “ECM organization“, cluster 2 was the only cluster with enriched “hydrogen peroxide (H_2_O_2_) catabolic/metabolic processes”. H_2_O_2_ has several and dose-dependent roles during wound healing. While disinfecting the wound tissue at relatively high concentrations, it promotes secretion of cytokines to improve dermal wound healing at lower concentrations [87, 88]. Cluster 2 fibroblasts expressed high levels of *gstp1*, a member of the family of glutathione transferases which in other instances have been demonstrated to protect cells for example from lipid peroxidation [89], as well as *hbba1* (S10 Fig.), a subunit of hemoglobin, which is known to attenuate H_2_O_2_-induced oxidative stress, intracellular glutathione depletion and cell death [90], thereby possibly alleviating negative effects of H_2_O_2_.

### The dynamics of ECM build up and breakdown

From our histology studies, we know that between 2 dpw and 4 dpw, the granulation tissue has in net grown, whereas between 4 dpw and 6 dpw, it has in net shrunk again (Fig 1B-D). To analyze the impact of fibroblast-expressed genes on ECM build up and breakdown in a systematic and integrative manner, we used the Matrisome AnalyzeR tool [91] to annotate genes encoding ECM molecules, quantifying overall expression levels of selected genes in selected cell populations. We created two main categories: ECM build up, which comprises the categories of ECM-affiliated proteins, secreted factors, proteoglycans, collagens, and ECM glycoproteins; and ECM breakdown, which comprises ECM regulators (see Materials & Methods). Both ECM build up and ECM breakdown-related genes were upregulated upon wounding. However, in contrast to the actual increase of granulation tissue sizes from 2 dpw to 4 dpw, and its decrease from 4 dpw to 6 dpw, expression of build-up genes, if at all, continued to increase across all later time points, while expression of ECM breakdown even started to decrease after 4 dpw. This was not only so when integrating over the entire fibroblast population (Fig 3D,E), but also for the individual fibroblast subclusters, with cluster 0 expressing most of these genes (S11 Fig.).

That the dynamics of ECM build-up and breakdown scores of fibroblasts do not, in fact, match with the actual dynamics of granulation tissue formation and regression might indicate that other cell types such as innate immune cells have a major impact. For example, *mmp13a*, the major collagen type I collagenase - the zebrafish lacks MMP1 and MMP8, the two other primary collagen type I collagenases in human [92] - was only weakly expressed in fibroblasts, but strongly in neutrophils, while *timp2b*, encoding the major MMP13 tissue inhibitor [93], was strongly expressed in fibroblasts, neutrophils and macrophages, with macrophage levels dropping during phases of granulation tissue regression (4 dpw, 6 dpw; S12 Fig.), possibly in net allowing stronger extracellular collagen I degradation. In addition, as mentioned above (Fig 2G, S3 Fig., S5 Fig.), macrophage subcluster 0 expresses the Mrc1 receptor implicated in the phagocytosis and further intracellular degradation of MMP-predigested collagens [55].

Yet, Matrisome AnalyzeR analyses integrating fibroblast, macrophage and neutrophil clusters yielded results similar to those for the fibroblasts only, with ECM degradation / ECM build-up ratios during granulation tissue regression (6 dpw) not being higher than during granulation tissue formation (2dpw; S11 Fig.). Together, this suggests that ECM-forming and ECM-degrading processes largely co-exist during all stages of wound healing and that comparably moderate changes in the balance between the two - with little impact on the overall program assessed by Matrisome AnalyzeR – might in net gradually push the granulation tissue from growth to regression.

### Molecular interactions between macrophages and fibroblasts

In addition to directly contributing to ECM formation and degradation, innate immune cells mainly affect these processes by instructing fibroblasts accordingly. During the descriptions of the macrophage and fibroblast subclusters (see above), we have already referred to some examples of potential macrophage-fibroblast, macrophage-macrophage and/or fibroblast-fibroblast signaling processes, e.g. via IL1, TNFα and CXCL19/CXCR4. Other prominent signals known for decades to be involved in macrophage – fibroblast signaling during cutaneous wound healing in mammals are Fibroblast Growth Factors (FGFs), Platelet-Derived Growth Factor (PDGF), and Transforming Growth Factor beta (TGFβ) [94].

Previous studies in our laboratory showed that global blockage of FGF signaling leads to failed granulation tissue formation similar to the effects of immune suppression, suggesting that innate immune cells might promote granulation tissue formation by secreting FGFs to stimulate fibroblasts [4]. To confirm these findings also in our scRNA-seq data, we checked the expression of Fgf ligands and Fgf receptors in fibroblasts and macrophages. All five receptors were expressed by the dermal fibroblasts at different levels during wound healing (Fig 4D), while some of the secreted Fgfs were expressed by macrophages (Fig 4E). This indicates that FGF signaling from macrophages to fibroblasts might indeed lead to fibroblast activation and granulation tissue formation, as proposed [4]. However, FGF receptors were also expressed by macrophages (Fig 4F), and Fgf ligands by fibroblasts (Fig 4G), suggesting that fibroblasts might also stimulate each other and might talk back to macrophages via FGF signaling.

**Fig 4.**
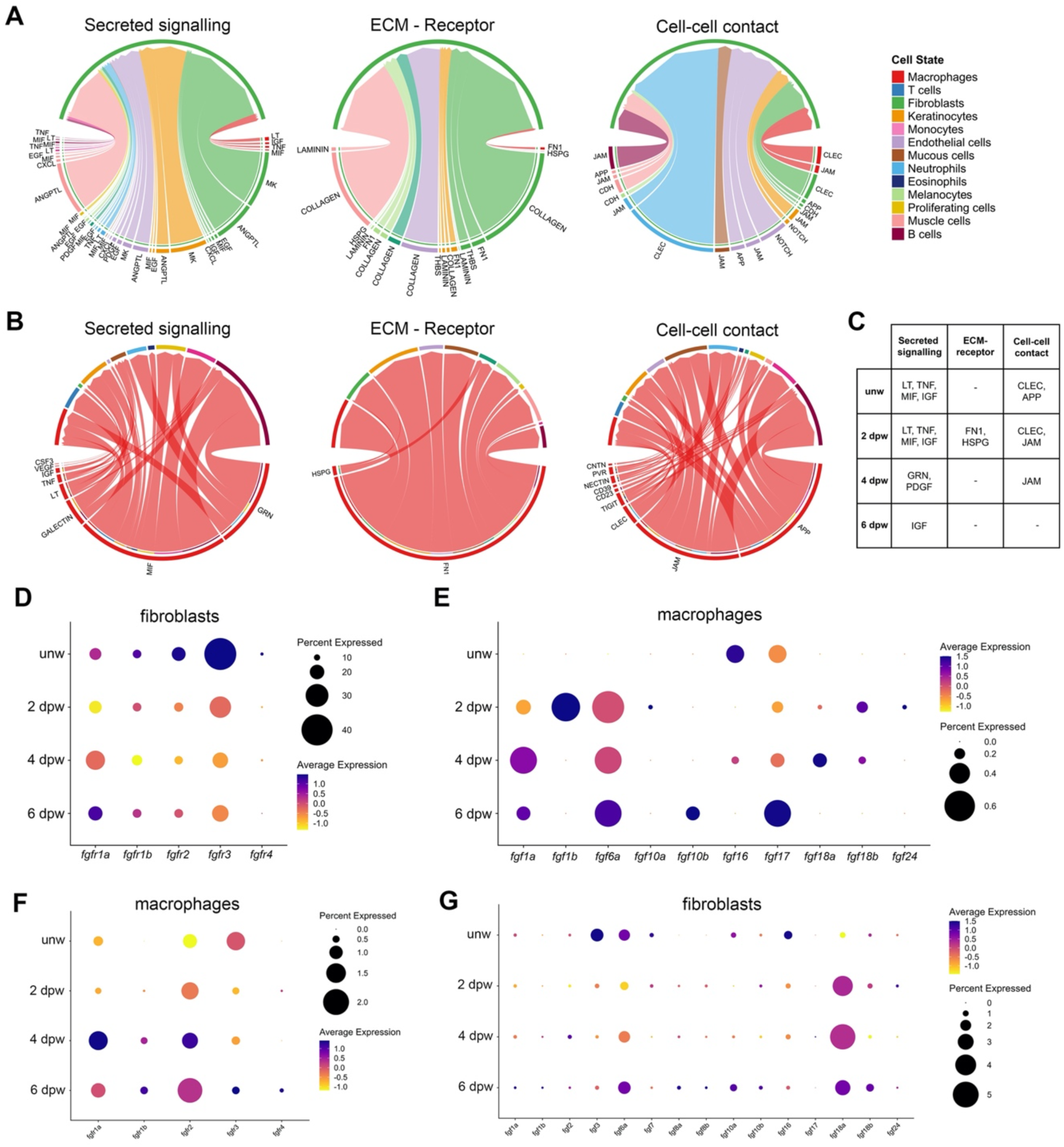
Ligand – receptor analysis between macrophages and fibroblasts. (A) Circos plots summarizing the active pathways of all time points from all cell clusters to fibroblasts. (B) Circos plots summarizing the active pathways of all time points from macrophages to all cell clusters. For corresponding Circos blots of specific time points, see Supplementary S8 Fig. (C) Summary table of the interactions between macrophages and fibroblasts across the different time points. (D-G) Dot plots showing expression level of fgf receptors and secreted fgf ligands in macrophages and fibroblasts, as found in the scRNA-seq data.

Similar expression patterns pointing to cross talk between multiple cells types were also observed for PDGF and TGFβ signaling. Thus, expression of the PDGF ligands (*pdgfab*, *pdgfba*) was not only initiated in macrophages upon wounding, but also in neutrophils and in fibroblasts themselves, while PDGF receptors (*pdgfra*, *pdgfrb*) were primarily and constitutively expressed in fibroblasts (S13 Fig.). This suggests that PDGF signaling also occurs during wound healing in adult zebrafish, and that the described mitogenic as well as chemotactic effects on fibroblasts [95] might originate from different innate immune cells and might involve autocrine signaling among fibroblasts.

Similarly, the zebrafish TGFβ1 ligand *tgfb1a*, in addition to being induced in macrophages, was also constitutively expressed in fibroblasts themselves, while expression of its paralog *tgfb1b* was primarily induced in neutrophils. Furthermore, at least one of the TGFβ type 1 receptors (*tgfbr1a*, *tgfbr1b*) and the type 2 receptors (*tgfrb2a*, *tgfrb2b*) were induced or constitutively expressed in fibroblasts (and in macrophages and neutrophils) (S13 Fig.), suggesting that the repair/fibrosis-stimulating effect of TGFβ signaling might in addition to the well-established instruction of fibroblasts by macrophages also involve instructions by neutrophils and by fibroblasts or among different fibroblast subclusters themselves.

To explore the potential interactions between macrophages and fibroblasts more systematically, we also performed CellChat analysis of our scRNA-seq data. The ligand receptor analysis was divided into three categories: secreted signaling, ECM – receptor and cell – cell contact, and analyzed for the interaction between all cell types and fibroblasts (Fig 4A, S14 Fig.) and between macrophages and all cell types (Fig 4B, S14 Fig.). The majority of the interaction between macrophages and fibroblasts were via secreted signaling, which of note changed during the different phases of wound healing (Fig 4C, S14 Fig.). In unwounded and 2 dpw skin, the main signaling processes were Lymphotoxin (LT) signaling, widely implicated in mammalian fibrosis [96], TNF signaling (see above) [97], Macrophage migration Inhibitory

Factor (MIF) signaling, long known for its central regulatory role during mammalian of wound healing [98] and Insulin Growth Factor (IGF) signaling, known to be up-regulated in many mammalian fibrotic diseases, promoting fibroblast proliferation as well as ECM production [99]. At 4 dpw, this changed to dominant signaling via PDGF (see above) and progranulins (GRN), known mammalian inflammatory regulators with described pro-as well as anti-fibrotic effects [100], whereas at 6 dpw, during granulation tissue regression, only some remaining IGF signaling was annotated.

Of note, in addition to fibroblast <-> macrophage signaling, the Circos plots for most annotated signaling pathways also indicated signaling between neutrophils and fibroblasts as well as signaling among macrophages and among fibroblasts (Fig 4A,B, S14 Fig), in line with the findings for FGF, TGFβ and PDGF signaling, described above.

In addition to signaling, potential cell-ECM and cell-cell contacts between macrophages and fibroblasts were predicted, the latter mainly via C-type lectin receptors (CLECs) and junction adhesion molecules (JAMs), both described with positive and negative implications in cell phagocytosis [101, 102]. However, these predicted macrophage-fibroblast contacts were surpassed by the same type of cell-cell contacts between neutrophils and fibroblasts. For both macrophage-fibroblast and neutrophil-fibroblast contacts, numbers were highest at 2 dpw and but dropped down to or even below the levels in unwounded skin during later stages of wound healing (Figs. 4A,B, S14 Fig.).

### *plod2* is expressed by wound fibroblasts

One other crucial interaction between macrophages and fibroblasts formerly reported by our laboratories causing imperfect healing of cutaneous wound in mice involves the resistin-like molecule RELMα. It is encoded by the *Retnla* gene, produced and secreted by macrophages, and instructs fibroblasts to generate Lysylhydroxylase 2 (LH2) encoded by the *Plod2* gene. LH2 function in turn leads to tight DHLNL crosslinking of collagens and scar formation [39]. Interestingly, according to phylogenetic analyses *retnl* genes first show up at the level of Coelacanthiformes (Latimeria) and are absent in teleosts like the zebrafish [103], suggesting that during evolution they were first invented during the aquatic-terrestrial translocation of the ancestors of land-based vertebrates.

Nevertheless, we found *plod2* in our scRNA-seq analyses to be expressed in fibroblasts of unwounded skin, and to be strongly up-regulated upon wounding (Fig 5A), in particular in fibroblast subcluster 0 and (myofibroblast) subcluster 4 (Fig 5B), and at stages of granulation tissue formation (2 dpw, 4 dpw), while numbers of expressing cells and expression levels, in contrast to many other genes (Figs. S9, S10), dropped again during granulation tissue regression (6 dpw) (Fig 5A). Of note, zebrafish in addition to *plod2* has two other genes (*plod1a*, *plod3*) encoding lysyl hydroxylases, Lh1 and Lh3 [104]. However, according to our scRNA-seq data, *plod1a* and *plod3* expression levels were much less affected by wounding than that of *plod2* (Fig 5C), suggesting that the wounding-induced increase of collagen DHLNL crosslinking (see below) is largely caused by LH2. Furthermore, Lh2 has two splice variants, with the longer variant containing an additional exon (13A) of 63 base pairs that codes for the telopeptidyl lysyl hydroxylase activity crucial for DHLNL crosslink formation [104]. Strong *plod2* expression in zebrafish wounds was confirmed via *plod2* in situ hybridization at 4 dpw (Fig 5F,G), while semiquantitative and quantitative RT-PCR of wound biopsies with specific primers for the longer splice isoform revealed that the expression of this relevant *plod2* variant was approximately 50-fold upregulated at 4 dpw compared to unwounded skin (Fig 5D), and had decreased again at 6 dpw (Fig 5E).

**Fig 5.**
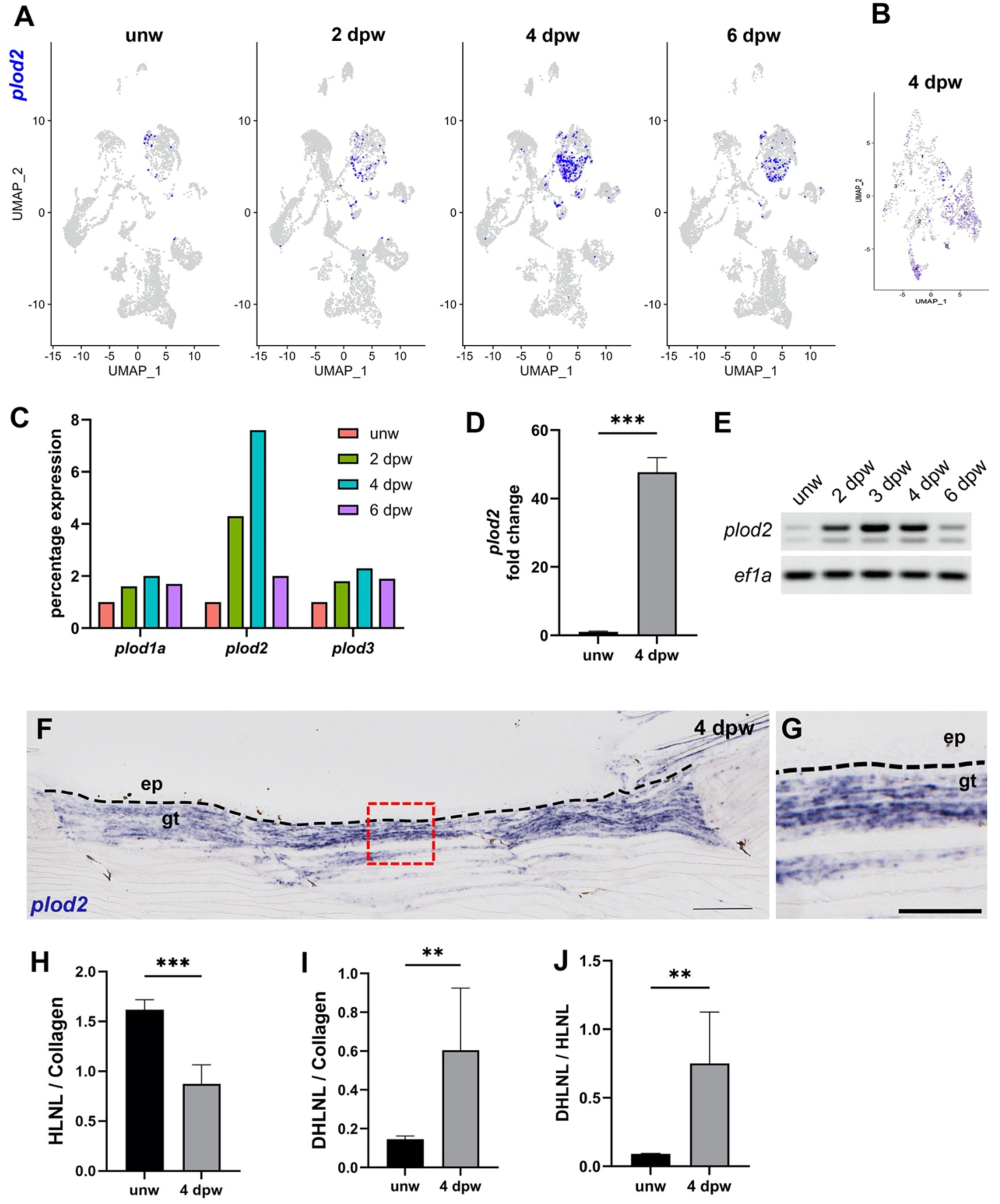
plod2 expression is upregulated in cutaneous wounds of zebrafish. (A) UMAP representation of plod2 expression across the time points. (B) UMAP representation of plod2 expression in fibroblasts at 4 dpw. (C) The expression levels of plod1a, plod2 and plod3 across the time points in scRNA sequencing data. (D, E) RT-PCR and qPCR results demonstrate that plod2 expression is upregulated upon wounding. Data represent mean ± SD, n = 3. (F, G) In situ hybridization results showing plod2 is expressed in the granulation tissue at 4 dpw. (H) The ratio of HLNL to collagen, (I) DHLNL to collagen and (J) DHLNL to HLNL crosslinks at 4 dpw. Data represent mean ± SD; n = 6 for unw; 2 for 4 dpw, with 3 pooled biopsies each. Numerical values for panels C, D, H, I and J can be found in S1 Data. Significances were determined with a two-tailed Student’s t test, *, P < 0.05; **, P < 0.01; ***, P < 0.001. Scale bars: 200 μm in F; 100 µm in G.

Since lysine residues in the telopeptide domain of collagens hydroxylated by LH2 will in subsequent steps give rise to DHLNL collagen crosslinks (rather than HLNL crosslinks when such residues remain non-hydroxylated) [36], we also biochemically analyzed DHLNL and HLNL crosslinks in wound and unwounded skin biopsy samples. Indeed, upregulation of *plod2* in zebrafish wounds coincided with a reduction in HLNL/collagen ratios (Fig 5H) and an increase in DHLNL/collagen ratios (Fig 5I), yielding an almost 5-fold increase in DHLNL/HLNL ratios at 4 dpw (∼0.15 to ∼0.75; Fig 5J). These numbers were similar to, and the changes upon wounding even stronger than those reported for mouse wounds (∼0.3 to ∼1.0) [39], indicating that in cutaneous zebrafish wounds, collagen crosslinking is as strong as in mouse wounds.

### *plod2* expression is stimulated by TGFβ signaling

Rather than by RELMα as in cutaneous wounds, *PLOD2* expression has in multiple other mammalian contexts been reported to be stimulated by TGFβ1/SMAD3 signaling [105–107]. Indeed, the promotor of the human *PLOD2* gene contains binding sites for the TGFβ1-activated transcription factor SMAD3, and binding of SMAD3 to this promotor element is required for TGFβ1-induced *PLOD2* expression in human fibroblasts, indicating that *PLOD2* is a direct TGFβ1/SMAD3 target gene [105]. To test whether in the absence of RELMα (see above), *plod2* expression in zebrafish cutaneous wounds might also be induced by TGFβ signaling, wounded fish were treated with the TGFβR inhibitor SB431542 between 3 dpw and 4 dpw.

qPCR analysis revealed that at 4 dpw, *plod2* mRNA levels were 5-fold lower in samples of SB431542-treated fish than in the control group (Fig 6A), indicating that wounding-induced *plod2* expression depends on and occurs via TGFβ signaling. Consistently, according to our scRNA-seq data, both TGFβ1-encoding zebrafish genes, *tgfb1a* and *tgfb1b*, were upregulated upon wounding at 2 dpw, especially in neutrophils and macrophages (Fig 6B), pointing to these innate immune cells as a crucial source of TGFβ1 to stimulate fibroblasts. Yet, fibroblasts might in addition, and particular at later stages, also stimulate each other via *tgfb1a* expressed at increased levels in wound fibroblasts at 4 dpw (see above; S13 Fig.). That wound fibroblasts receive and transmit such TGFβ signals is indicated by two lines of evidence. First, our scRNA-seq analysis revealed that expression of *tgfbi* (transforming growth factor beta induced), a known downstream target and marker for TGFβ signaling [108], was upregulated in fibroblasts (and in macrophages) at 4 dpw, and in the same fibroblast subclusters as *plod2* (Fig 6C). Second, immunostainings of phosphorylated Smad3 in a transgenic line with fluorescently labelled fibroblasts demonstrated the presence of activated Smad3 in the nuclei of wound fibroblasts at 4 dpw (Fig 6D). Taken together, the wound dermal fibroblasts in zebrafish also express *plod2* as they do in mammals, and this expression is dependent on TGFβ signaling, in contrast to mammalian dependency on RELMα.

**Fig 6.**
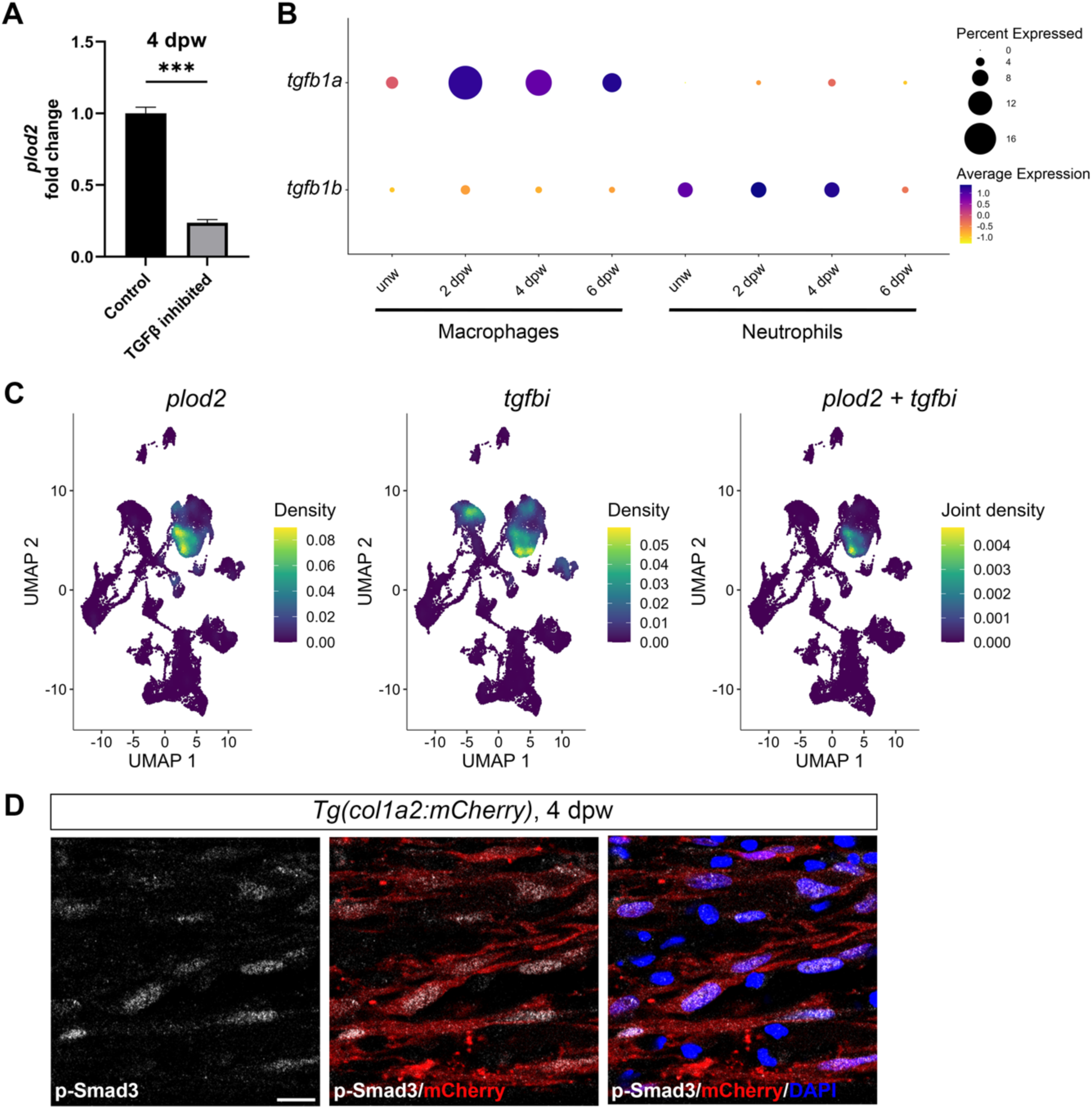
*plod2* expression is downstream of TGFβ1 signaling. (A) qPCR results showing plod2 expression is downregulated at 4 dpw upon inhibition of TGFβ signaling via TGFβR inhibitor (SB431542). Data represent mean ± SD, n = 3. Numerical values for panel A can be found in S1 Data. (B) Dotplot showing the expression of tgfb1 paralogs in macrophages and neutrophils upon wounding. (C) UMAP representation of expression density for plod2 and tgfbi and co-expression of both transcripts in scRNA sequencing data. (D) Immunostaining of phospho-Smad3 and mCherry at 4 dpw sections of the transgenic fibroblast reporter line. Significances were determined with a two-tailed Student’s t test, *, P < 0.05; **, P < 0.01; ***, P < 0.001. Scale bars: 10 μm in D.

### Gain or loss of Lh2 function does not compromise regenerative ability

Reduced *Plod2* expression levels caused by loss of RELMα leads in cutaneous mouse wounds coincides with reduced wound scarring. Similarly, direct siRNA-mediated knockdown of Plod2 alleviates liver necrosis by making collagen fibers deposited by liver myofibroblast aligned in a more orderly manner and more easily degradable [109], pointing to a pro-fibrotic effect of Plod2. Yet, despite highly up-regulated *plod2* expression levels, zebrafish wounds eventually heal perfectly, with a full regression of the transiently formed granulation tissue. Given that in contrast to most other fibroblast markers, expression of *plod2* drops again during granulation tissue regression (see above; Fig 5A,D), we investigated whether Plod2 might at least affect the dynamics or degree of granulation tissue formation and regression, performing gain- and loss-of-function experiments. For temporally controlled gain of function, a transgenic zebrafish line was generated carrying a heat shock inducible promoter driving the expression of *plod2*, *Tg(hsp70l:plod2-p2A-EGFP)*. Histological analysis of tissue sections from repetitively heat-shocked transgenic and non-transgenic control fish at 4 dpw, when granulation tissue is normally maximal, and at 8 dpw and 12 dpw, when granulation tissue has regressed, revealed that despite a several-fold increase of *plod2* expression in heat-shocked transgenic fish (Fig 7A), the sizes of granulation tissue were not changed upon forced *plod2* overexpression compared to heat-shocked non-transgenic siblings (Fig 7B,C). Collagen I immunofluorescence and Transmission electron microscopy (TEM) further revealed that the amounts of fibrillar collagen type I (Fig 7D,E, S15 Fig.), as well as the diameters and the alignment of collagen fibers (Fig 7F) within the granulation tissue were not affected by forced *plod2* overexpression compared to wild-type siblings.

**Fig 7.**
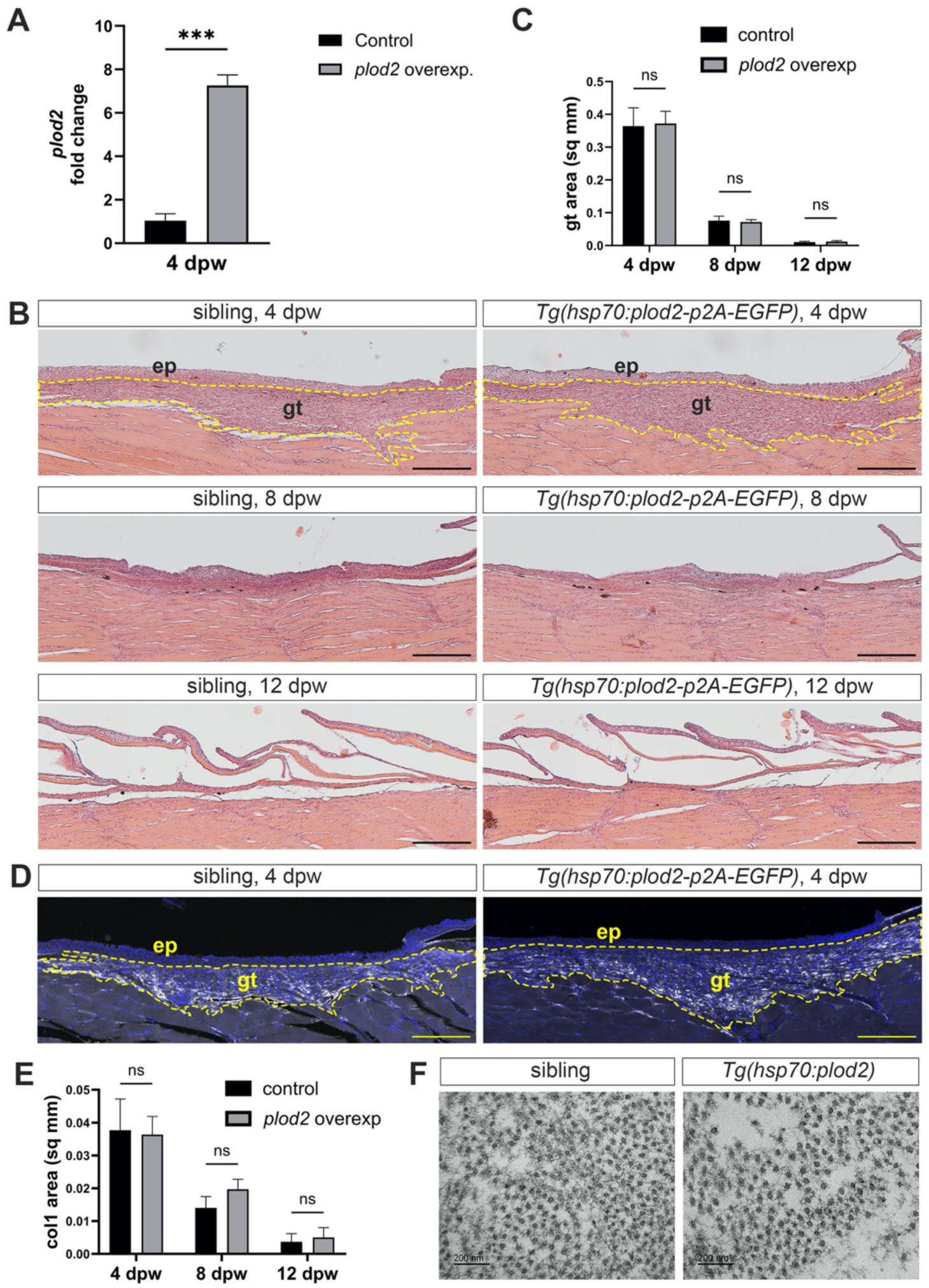
Gain of Lh2 function does not compromise granulation tissue resolution. (A) qPCR results showing that in from 0 – 4 dpw daily heat-shocked Tg(hsp70l:plod2-p2A-EGFP) fish, plod2 expression is over sevenfold upregulated at 4 dpw compared to heat-shocked non-transgenic siblings. Data represent mean ± SD, n = 3. (B) Representative H&E stainings on sections of daily heat-shocked Tg(hsp70l:plod2-p2A-EGFP) fish and their control siblings at 4 dpw, 8 dpw and 12 dpw, respectively (yellow dashed lines mark the granulation tissue). (C) Quantification of granulation tissue sizes in sections as shown in (B), revealing that sizes were not changed upon forced plod2 overexpression compared to wild-type siblings. Data represent mean ± SD, n = 3. (D) Representative images of immunostaining against collagen I protein (gray) at 4 dpw on paraffin sections, counterstained with DAPI (blue); for other stages, see Fig. S15. (E) Quantification of the collagen I stained areas at 4 dpw, 8 dpw and 12 dpw showing no changes upon forced plod2 overexpression compared to wild-type siblings. Data represent mean ± SD, n = 3. Significances were determined with a two-tailed Student’s t test, *, P < 0.05; **, P < 0.01; ***, P < 0.001. (F) TEM analysis of Tg(hsp70l:plod2-p2A-EGFP) fish, demonstrating that the diameters and alignment of collagen fibres were not changed at 4 dpw compared to wild-type siblings. Numerical values for panels A, C and E can be found in S1 Data. Scale bars: 200 μm in A-C, F; 200 nm in E. ep: epidermis, gt: granulation tissue.

For loss of function, we carried out the same type of analyses with a formerly isolated *plod2* mutant carrying a non-sense mutation that results in a C-terminally truncated (679 instead of 754 amino acid residues), non-functional protein [110]. Although displaying the described skeletal defects pointing to a requirement of LH2 for proper collagen I fiber organization in bones and for bone stability, cutaneous wound healing was not affected in the mutants, with no significant difference in granulation tissue sizes compared to wild type siblings (A,B in S16 Fig.), normal intensities and patterns of granulation tissue collagen I in immunostainings (C,D in S16 Fig.), and no differences in the diameters and alignments of granulation tissue collagen fibers in TEM images (E in S16 Fig.). Taken together, the overall regenerative ability was not compromised during cutaneous wound healing in zebrafish upon forced gain or loss of Lh2 function, indicating that even higher or lower levels of this pro-fibrotic factor do not affect the dynamics of granulation tissue formation and regression.

## Discussion

Cutaneous wound healing represents a complex biological process which involves the interaction of diverse cell types and signaling pathways [5]. In contrast to fibrotic scarring observed in mammals, zebrafish possess the capability to perfectly restore lost or damaged tissue to its original state without scarring, although wounds also display “temporary fibrosis” in a granulation tissue that fully regresses during later stages of wound healing [4]. The same applies to other perfectly regenerating organs of adult zebrafish, for example the heart after cryoinjury, which also displays temporary formation of an excessive scar that later regresses [111]. In order to dissect the cellular and molecular processes underlying granulation tissue formation versus regression, we performed scRNA-seq for such different stages of cutaneous wound healing in adult zebrafish.

### Cellular and molecular dynamics and heterogeneities during zebrafish cutaneous wound healing

During mammalian skin wound healing, the wound microenvironment undergoes dynamic changes, with different cell types participating in tissue repair. Particularly, macrophages and fibroblasts are key players during this process [11, 112], and this also seems to be so during zebrafish wound healing. Macrophages arrive in the wound before fibroblasts and later start to drop in numbers earlier than fibroblasts [4]. Yet, both cell types co-exist and cooperate not only during the formation, but also during the regression of granulation tissue.

As in mammals, macrophages and fibroblasts of zebrafish skin wounds exhibited a whole spectrum of activation states with unique gene expression profiles defining multiple subclusters each - although only one of our annotated fibroblast subclusters was wound-specific, compared to four of seven in cutaneous mouse wounds [24]. Macrophage subclusters with preferential pro-inflammatory, anti-inflammatory and/or pro-regenerative characteristics were identified. However, most of these subclusters co-existed during all stages of zebrafish wound healing, suggesting that pro- and anti-inflammatory processes largely overlap in time - although our calculated overall inflammation score clearly peaks at early stages of wound healing (Fig 2). A similar co-existence and temporal overlap during granulation tissue formation and regression was observed for most fibroblast subclusters, the main producers of the ECM (Fig 3). In addition to fibroblasts, innate immune cells also seem to directly contribute to ECM production - as also recently shown for mammalian wounds [60] - and ECM degradation. For example, macrophage subcluster 5 showed strong expression of genes encoding fibrillar collagens as well as basement membrane components (S5 Fig.), while neutrophils showed strongest expression of *mmp13b*, together with *mmp13a* the only “professional” collagenase encoded by the zebrafish genome (S12 Fig.). Of note, however, even when integrating fibroblasts, macrophages and neutrophils, our ECM build up / ECM breakdown analyses yielded at first sight puzzling results, with formation/degradation ratios during granulation tissue formation not being higher than during granulation tissue regression (S11 Fig.). Together, this points to a large degree of co-existence of ECM-forming and ECM-degrading processes during all stages of wound healing. In addition, it suggests that rather specific temporal changes in particular cell types and/or in the expression levels of rather few and particularly important factors might make the difference between net formation or net regression of granulation tissue.

Of note, in addition to changes in cells and potential regulatory genes, our scRNA-seq data also point to dynamic changes within the ECM itself. Thus, while Col I, the main fibrillar dermal collagen, was expressed at rather unaltered levels during all stages of wound healing, Col V, the other fibrillar dermal collagen, as well as Col II and Col X, fibrillar and network-forming collagens mainly present in (hypertrophic) cartilage, respectively, were downregulated during granulation tissue formation compared to unwounded skin, but strongly upregulated during granulation tissue regression. In contrast, Col XII, a FACIT associated with Col I-containing fibrils, Col XI, the other fibrillar collagen mainly present in cartilage, and Col XVIII were upregulated both during granulation tissue formation and regression (Fig 3G, S9 Fig.). A strong upregulation particularly during granulation tissue regression was also observed for other ECM proteins, such as Fibronectin (*fn1b*), the Cartilage Intermediate Layer Protein 1 (*cilp1*) and the small leucine-rich proteoglycan Osteoglycin (*ogna*) (S10 Fig.). Together, this points to crucial differences in the ECM composition of the growing versus the regressing granulation tissue, with the regressing ECM displaying striking similarities to cartilage ECM (Col II, Col X, Col XI, Cilp1, Ogna).

### Myofibroblasts during zebrafish cutaneous wound healing

In mammals, myofibroblasts are considered as “the primary ECM-producing cell types executing fibrosis” [113], including scarring of skin wounds [23]. In addition, their contractile function contributes to the closure of mammalian skin wounds [114]. For the closure of zebrafish wounds, myofibroblasts are most likely dispensable, as the wounds close within hours and even before fibroblasts have entered the wound bed [4], purely driven by re-epithelialization movements of the epidermis [40]. Accordingly, we had thus far failed to detect myofibroblasts in zebrafish skin wounds with anti-smooth muscle actin antibodies, although myofibroblast-like cells have described during zebrafish heart regeneration [111, 115, 116].

Interestingly, although not constituting a distinct fibroblast subcluster, we now identified skin wound fibroblasts that display a transcriptional profile similar to mammalian myofibroblasts. Of note, they are not present in unwounded skin and, compared to other fibroblast subtypes, show a stronger reduction in cell numbers during later stages of wound healing, when granulation tissue regresses (Fig 3, S8 Fig.). In light of the described strong dependence on the relative abundance of myofibroblasts on their beneficial tissue-remodeling versus detrimental pro-fibrotic effects [22, 23], it is tempting to speculate that it is this particular kinetics of myofibroblasts that contributes to temporary versus persistent fibrosis during zebrafish versus mammalian wound healing. Future studies have to address the particular role of myofibroblasts during zebrafish granulation tissue formation and the mechanisms underlying their disappearance during later stages of zebrafish wound healing.

### Other pro-fibrotic pathways and Lh2 during zebrafish cutaneous wound healing

In mammalian wounds, other pro-fibrotic genes and cell types beyond myofibroblasts have been shown to contribute to persistent fibrosis and scarring. Puzzlingly, according to our scRNA-seq analysis, most of them are also present in zebrafish skin wounds, for instance En1, a transcription factor marking mammalian fibroblasts that upon ablation or blockage of En1 activation prevent scarring of injured skin [27, 28], or Periostin (*postna*, *postnb*; S8 Fig.), a secreted protein made by fibroblasts that upon ablation alleviate scarring in injured hearts [70]. Future cell ablation and genetic gain- and loss-of-function experiments have to reveal the roles of these cell types and genes during cutaneous wound healing in zebrafish.

Another described mammalian pro-fibrotic factor, for which we have already here performed functional studies, is Lysylhydroxylase 2 (Lh2), encoded by the *Plod2* gene. In mammalian wounds, *Plod2* expression is induced in dermal fibroblasts by RELMα released by macrophages, leading to more stable DHLNL-dependent crosslinking between collagen fibers and persistent fibrosis [39]. Given that *retln* genes encoding RELMα and related factors are absent from fish genomes, we had speculated that zebrafish wounds might lack *plod2* expression and collagen DHLNL crosslinks, and that this has a crucial impact on granulation tissue regression. Surprisingly, however, our results demonstrated that *plod2* expression and collagen DHLNL crosslinks are also induced in zebrafish wounds, in this case induced by TGFβ signaling, a well-known mediator of mammalian fibrosis in other pathways and contexts (Fig 5). Moreover, Lh2 gain or loss of function did not compromise the regenerative ability of zebrafish (Fig 6), challenging this particular conception of fibrosis regulation [39], and pointing to the existence of alternative mechanisms allowing granulation tissue regression, ECM breakdown and thereby scar-free wound healing even in the presence of those more stable collagen fibrils.

### Implications for wound healing therapies

The long-term goal of our work is the identification of specific cell types or genes making the difference between perfect healing with only temporary fibrosis in fish versus imperfect healing with persistent fibrosis and scarring in mammals, including human. In principle, we would expect that fish wounds contain anti-fibrotic factors absent in mouse wounds, and/or that mouse wounds contain pro-fibrotic factors absent in fish wounds. In this light, RELMα described above was a good candidate for the second category, which unfortunately turned out negative, as its absence in fish can be compensated by TGFβ. This makes it quite likely that similar compensatory mechanisms could also be activated when targeting RELMα in human skins to alleviate scarring. Thus, concomitant targeting of RELMα and TGFβ signaling might be a reasonable approach. Candidates for anti-fibrotic factors absent in mouse wounds were not so apparent in our analyses. They might be among the genes with no annotated function as yet that were strongly up-regulated in zebrafish wounds, for instance si:dkey-21e2.8, which is strongly expressed in fibroblast subcluster 3 of fish wounds; S7 Fig.). Yet, also our own BLAST searches did not unravel any orthologs in vertebrate classes other than fish, thus it remains unclear whether it is a least part of a pathway shared between fish and mammals, and how to possibly pharmacologically promote this pathway in human wounds. Alternatively, rather mild differences among pro- and anti-fibrotic factors shared by fish and mammalian wounds might underlie perfect versus imperfect wound healing in fish versus mammals. Consistent with this possibility, we found most factors with described pro-fibrotic effects during mammalian wound healing also present in zebrafish wounds (see above). On one hand, it is challenging to identify such most likely multi-factorial differences in cross-species scRNA-seq analyses [10, 45, 51, 54] [117]. On the other hand, however, when successful, it might be instrumental to develop novel or modified therapeutic interventions in wound healing-related human disorders, possibly using a combination of already established pharmacological agonists and antagonists.

In conclusion, our findings set the base for further and more systematic cross-species scRNA-seq analyses to elucidate crucial differences in the cellular and molecular dynamics during the different phases of perfect cutaneous wound healing in fish versus fibrotic wound healing in mammals, with the ultimate goal to develop new preventative and reparative therapies for skin repair in human.

## Materials and Methods

### Zebrafish lines and wounding

Six to eight months of age adult zebrafish Tüpfel long fin (TL) and Ekkwill (EK) wild type strains were used. Transgenic and mutant lines used in this study were: *Tg(col1a2:mCherry-NTR)*^cn11Tg^ [118], *plod2*^sa1768^ mutants [119]. Transgenic line *Tg(hsp70l:plod2-p2A-egfp)*^fr62Tg^ was generated in this study. Full-thickness wounds of adult fish with a diameter of ∼2 mm were introduced with a clinical dermatology laser as described [4].

Zebrafish were raised at 28°C on 14 hours/10 hours light/dark cycle and fed with paramecia, dry flake food and live/frozen artemia daily. Feeding of larvae and adult fish and monitoring of the quality of the system water were done by the institute’s animal care takers daily.

The zebrafish wound healing experiments were approved by the local and federal animal care committees (City of Cologne: 8.87-50.10.31.08.129; LANUV Nordrhein-Westfalen: 81-02.04.2018.A097, 81-02.04.2022.A104, 81-02.04.40.2022.VG005).

### Genotyping of *plod2* mutants

*plod2*^sa1768^ mutants were genotyped using PCR amplification with the primers listed in Table 1. This yields a 264 bp amplification product which upon digestion with *Mbo*II enzyme (New England Biolabs) yields bands of 153 bp, 70 bp and 41 bp in wild type genomic DNA, but bands of 223 bp and 41 bp in mutant genomic DNA.

**Table 1.**
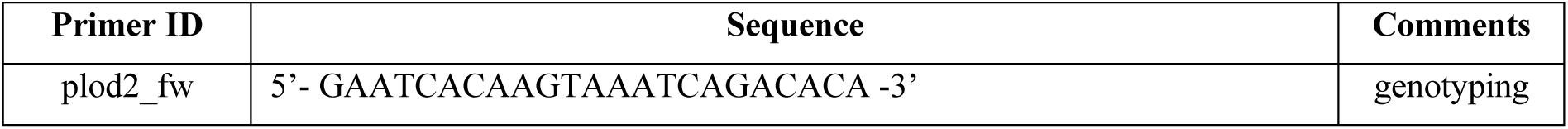
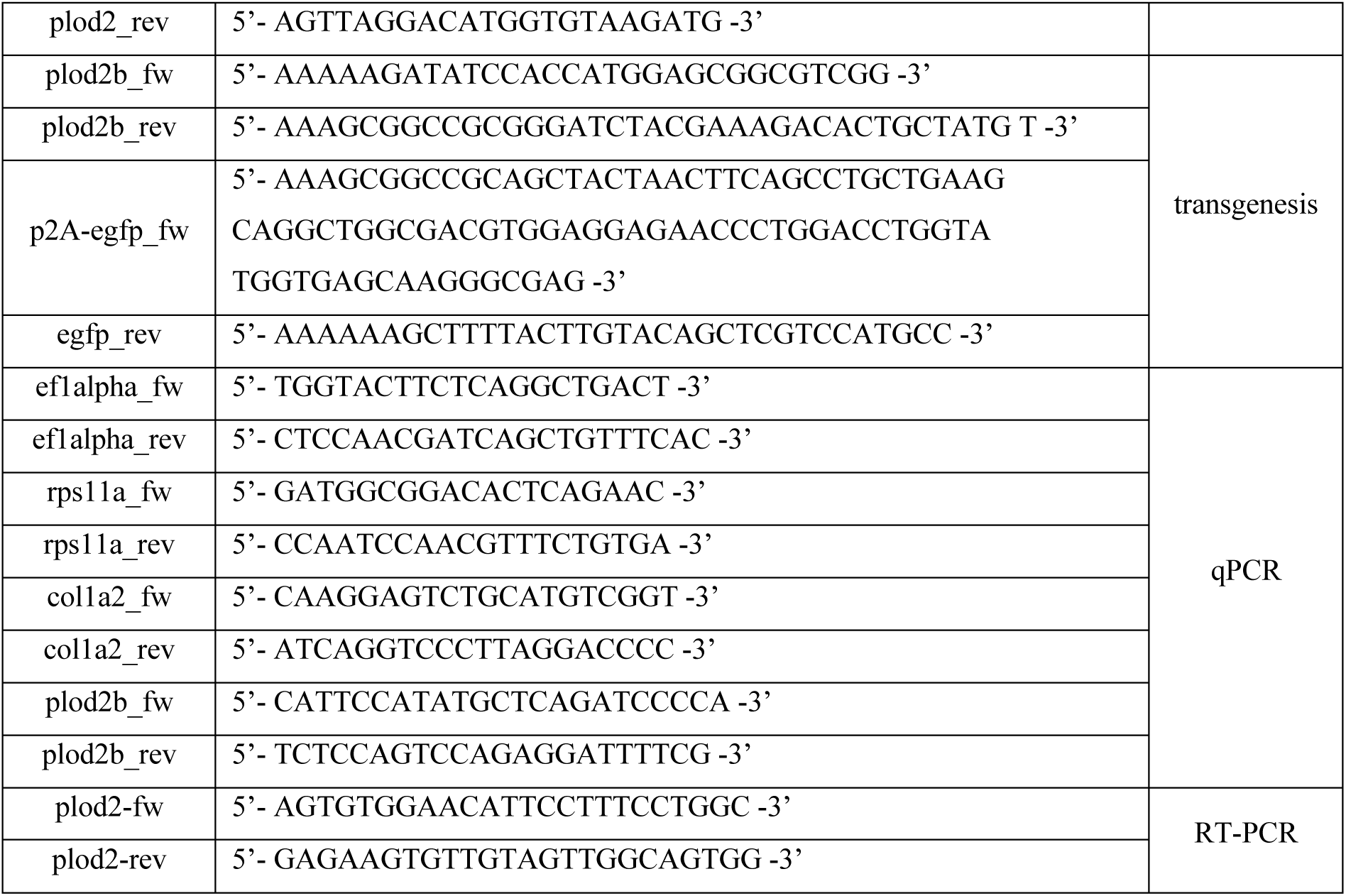
Primer sequences used in this study.

### Plasmid construction and transgenesis

For cloning of the construct hsp70l:plod2-p2A-egfp, the hsp70l promoter region was digested with *Xba*I and *Eco*RV restriction enzymes from a pBSIISK-hsp70l plasmid. Zebrafish *plod2* ORF was amplified using primers listed in Table 1 and digested with *Eco*RV and *Not*I restriction enzymes. The egfp-polyA part was amplified from p3E-egfp-ployA plasmid (Tol2 kit; {Kwan, 2007 #169}), with the p2A part added via the primers during amplification, followed by digestion of the PCR product with *Not*I and *Hind*III restriction enzymes. Finally, pminiTol2 plasmid was digested with XbaI and HindIII restriction enzymes and ligation was done between linearized pminiTol2 plasmid backbone and all three inserts together. The final construct was checked with Sanger sequencing.

The final plasmid construct was microinjected into fertilized 1-cell-stage embryos using micro-manipulator. The final concentrations of the constructs in the injection mix were 25 ng/μl together with 25 ng/μl Tol2 mRNA and 5 nl of the solution mix was injected inside of the cell. Embryos were raised and transgene expression was checked with fluorescent markers.

### Tissue labelling procedures

For paraffin embedding, samples were fixed in 4% paraformaldehyde/phosphate buffered saline (PBS) overnight at room temperature, decalcified in 0.5M EDTA (pH 8.0) for 5 days, dehydrated in a graded series of alcohols, cleared in Roti-Histol (Carl Roth, Karlsruhe, Germany), and embedded in paraffin. For cryosectioning, the trunk of the fish was cut in smaller pieces and fixed with 4% PFA/PBS overnight at 4°C. Fixed samples were washed with PBS, decalcified in 0.5M EDTA (pH 8.0) for 3 days at 4°C. Afterwards, the samples were incubated with 30% sucrose/PBS solution for two days at 4°C before embedding in OCT embedding matrix (Carl Roth, Germany). Both paraffin and OCT embedded blocks were sectioned at 10 μm thickness.

Immunofluorescence and histological analyses were performed using standard protocols. Antigen retrieval step was performed additionally for immunofluorescence analysis with 10 mM citrate buffer (pH 6.0) for 15 minutes. Stainings were performed with the primary antibodies mouse anti-mCherry (1:400, Abcam, ab125096), rabbit anti-collagen I (1:200, Abcam, ab23730), rabbit anti-phospho-Smad3 (1:400, Abcam, ab52903).

In situ hybridization on cryosections was performed as previously described [120]. Briefly, tissue sections were fixed again 5 min with 4% PFA, treated with proteinase K (5 μg/ml, 30 min) and PBS washes. Slides were incubated in triethanolamine-acetic anhydrate pH 8.0 5 min, then dehydrated in a series of EtOH. Sections were incubated for 3 h with hybridization solution (50% formamide, 5X SSC, 1X Denhardt, 10% dextran sulfate, 1 mg/ml tRNA). Digoxigenin-labeled *plod2* probe was generated by in vitro transcription [104]. Slides were blocked with 10% sheep serum in PBST at room temperature for 3 h, incubated overnight at 4°C with alkaline phosphatase-labeled anti-DIG antibody (Roche) and enzymatic activity was detected with BCIP and NBT.

### Drug treatments and heat shock induction

TGFβR inhibitor (SB431542, Sigma-Aldrich, S4317)) was administered to adult zebrafish according to the indicated times in E2 medium with 50 μM final concentration. All drug treatments were done with a final concentration of 0.1% DMSO. Drug-containing E2 media were changed every 24 hours as necessary.

Heat induction for adult zebrafish was done by transferring them to pre-heated 38°C system water. Adult fish were kept in a 38°C incubator for 2 hours before transferring them back to the system.

### qPCR analysis

Adult zebrafish wound tissue samples were collected using a biopsy punch with a diameter of 4 mm. Collected tissues for total RNA extraction were snap frozen with dry ice before adding Trizol reagent. Tissue samples were homogenized and kept at −80°C for at least overnight until total RNA extraction was done according to the Monarch Total RNA Miniprep Kit protocol. cDNA synthesis was done with 1 μg of RNA with the iScript cDNA Synthesis Kit. The qPCR for the gene expression analysis was performed using the SYBR Green PCR Master Mix with the primers listed in Table 1. Fold change was calculated using the ΔΔC_T_ method.

### Transmission electron microscopy

For TEM analysis, the trunk of adult zebrafish was cut in smaller pieces and fixed overnight at 4°C with 2% PFA + 2% GA solution. Fixed samples were washed with PBS, decalcified in 0.5 M EDTA (pH 8.0) for 3 days at 4°C. Afterwards, samples of 1-2 mm thickness were incubated 0.1M cacodylic acid and re-fixed with 2% osmium tetroxide (Science Services) in 0.1M cacodylic acid, and washed 4 times for 15 min in 0.1M cacodylic acid. Through a series of ethanol 50-100%, a mixture ethanol/ propylene oxide and 100% propylene oxide, the tissue was then embedded in EPON (Science Services). After embedding, the tissue was cut in 70 nm sections with a ultramicrotome (UC6, Leica) on a grid. Grids with sections were contrasted with 1.5% uranyl acetate aqueous solution for 15 min at 37°C, incubated for 4 min in lead citrate solution (Leica Ultrastain II), washed again five times in water and dried on a filter paper.

Images were acquired with a transmission electron microscope (JEM 2100 Plus, JEOL), a OneView 4K camera (Gatan) with DigitalMicrograph software at 80 KV at room temperature.

### Crosslink analysis

Collagen crosslink analysis of uninjured skin and three pooled biopsies of cutaneous wounds per stage was performed as previously described [39, 121]. Briefly, tissue specimens were reduced, denatured, and digested, and analysis was performed with an amino acid analyzer (Biochrom 20).

### Preparation of single cell suspensions

For single cell RNA sequencing, single cell suspensions were prepared. For this task, unwounded skin tissue from three adult zebrafish was collected by scraping off the skin and pooling in PBS (Mg^2+^ and Ca^2+^-free). In addition, the wound tissues for 2 dpw, 4 dpw and 6 dpw were collected using a biopsy punch with a diameter of 4 mm from three adult zebrafish each. PBS was replaced with 0.25 mg/ml Liberase TM in PBS solution and the tissues were incubated at 30°C for an hour with slow agitation (∼300 rpm). The solution was pipetted every 10-15 minutes. After successful digestion of the tissue, the solution was mixed with 5% FBS/PBS and filtered through a cell strainer with 70 μm nylon mesh and then a cell strainer with 40 μm nylon mesh in 50 ml Falcon tubes. The falcon tubes were centrifuged at 500 x g for 5 minutes and the pellet was resuspended with 2% FBS/PBS. The viability of the cells were more than ninety percent. The single cell suspensions were kept on ice until they were processed at the Cologne Center for Genomics, University of Cologne. 10.000 cells per condition was aimed for single cell RNA sequencing.

### Clustering of Single Cell Data Matrix

Single cell suspensions were processed, sequenced and single cell library construction was mapped to the zebrafish transcriptome by Cologne Center for Genomics, University of Cologne using 10x Genomics 3’scRNA seq application (Version 3’v.3.1) with Cell Ranger software used to generate feature-barcode matrices. Single cell RNA sequencing data was analyzed using RStudio program with Seurat (V4.3.0) software package for R using standard quality control [42, 122–125], normalization, integration and analysis steps. Low quality cells were excluded from downstream analysis according to following criteria for retaining cells: 500 < nUMI < 30.000, 200 < nGene < 4500, mito < 25%. The top 30 most differentially expressed genes were identified for each cluster and the most likely cell type was annotated for each cluster according to expression patterns of the genes found in public databases.

### Gene Ontology (GO) enrichment analysis

To analyze and visualize functional profiles of differentially expressed gene clusters across cell subtypes, we applied clusterProfiler R package [126] as indicated in accessible vignette (https://yulab-smu.github.io/clusterProfiler-book/). GO enrichment analysis was performed using the list of significantly upregulated genes. GO terms with adjusted p-value < 0.05 were considered significantly enriched.

### STRING Analysis

Overexpressed genes in the macrophage population of the 2 dpw sample (261 genes) were compared with the unwounded sample. The gene list was queried in STRING (version 11.5) using the Multiple Proteins by Names/Identifiers function. Functional enrichment analysis was performed under Biological Process (Gene Ontology) to identify pathways associated with inflammation. Among the enriched pathways, “GO:0006954 – Inflammatory response” was identified.

### Extracellular matrix (ECM) analysis

We used Matrisome AnalyzeR package to characterize ECM composition at single-cell resolution [91]. Differential gene expression analysis was first performed by comparing each sample to the unwounded sample. Genes significantly overexpressed in each sample were selected for downstream ECM-specific annotation following scRNA-seq analysis for Seurat objects provided workflow (https://github.com/Matrisome/MatrisomeAnalyzeR).

### Cell–Cell communication analysis

CellChat R package [127] was employed to determine potential ligand-receptor communication in our scRNA-seq gene expression data by using CellChatDB.zebrafish, according to the available tutorial (https://github.com/sqjin/CellChat).

## Quantification and statistical analysis

For the measurement of the granulation tissue areas, FIJI-ImageJ software was used and the area was drawn manually. Statistical analysis was performed using GraphPad Prism software. A two-tailed Student’s t test was performed for comparison of two groups. Significances were: *, P < 0.05; **, P < 0.01; ***, P < 0.001.

## Ethics Statement

The zebrafish studies were approved by the animal welfare office of the state of Northrhine-Westfalia (LANUV), approval numbers: 81-02.04.2018.A097, 81-02.04.2022.A104, 81-02.04.40.2022.VG005.

## Data availability

The single-cell RNA-sequencing data (GSE306419) used in this study can be accessed in the public database Gene Expression Omnibus (https://www.ncbi.nlm.nih.gov/geo/).

## Acknowledgements

We thank Andy Willaert from Ghent University, Belgium, for the *plod2*^sa1768^, Nadia Mercader Huber from the University of Bern, Switzerland, for the *tg(col1a2:mCherry-NTR)*^cn11Tg^ zebrafish lines, Beatrix Martini from the CECAD Imaging Facility for her help with the TEM analysis, the Cologne Center for Genomics for their technical assistance in performing the single-cell RNA sequencing, and Joy Armistead for proofreading of the manuscript. Work in the laboratory of MH was supported by the German Research Foundation (DFG; Research Grant project No. 453407124). FJMM thanks the Margarita Salas Foundation, Spain, for his long-term postdoctoral fellowship, IK thanks Sabine Werner for her support.

## Author contributions

Conceptualization: IK, SE, MH; Data curation: IK, MM, PC, JB; Formal Analysis: IK, FJMM, KH, NR; Supervision: SE, MH. Writing – original draft: IK, MH; Writing - review and editing: SE, MH.

## Supporting Information

**S1 Fig.** Cellular composition of zebrafish cutaneous wounds. UMAP representation of single cell RNA sequencing data after integration of four datasets and cell clustering results across the time points.

**S2 Fig.** Identification of different cell clusters during zebrafish wound healing. Heat map of the top 10 differentially enriched genes in each cell cluster.

**S3 Fig.** Identification of macrophage subclusters. Heat map of the top 10 differentially enriched genes in macrophage subclusters.

**S4 Fig.** The roles of macrophages during cutaneous wound healing. GO analysis of macrophage subclusters in unwounded skin and at 4 dpw and 6 dpw.

**S5 Fig:** UMAP representations of selected macrophage-specific genes across the different stages of wound healing (A) all clusters in unwounded skin (unw) and at 2 dpw, 4dpw, 6 dpw (B) macrophage cluster at 2 dpw *mrc1b, cxcl19, col1a1a, col4a1, col4a2, lama4, f13a1a, esr2a, cpn1, tnfa, tgfb1a, cxcr4a, cxcr4b*

**S6 Fig.** Identification of fibroblast subclusters. Heat map of the top 10 differentially enriched genes in fibroblast subclusters.

**S7 Fig.** The roles of fibroblasts during cutaneous wound healing. GO analysis of fibroblast subclusters in unwounded skin and at 2 dpw and 6 dpw.

**S8 Fig.** UMAP representations of myofibroblast-specific genes and marker genes for other fibroblast cell types implicated in fibrosis across different stages of wound healing (A) all clusters in unwounded skin (unw) and at 2 dpw, 4dpw, 6 dpw (B) fibroblast cluster at 4 dpw *acta2, ankrd1a, desma, cdh15, en1b, postna, postnb*

**S9 Fig.** UMAP representations of selected collagen-encoding genes across the different stages of wound healing (A) all clusters in unwounded skin (unw) and at 2 dpw, 4 dpw, 6 dpw (B) fibroblast cluster at 4dpw *col1a1, col1a2*, *col2a1*, *col5a2b*, *col12a1b*, *col11a1b*, *col10a1, col18a1b*, *col4a1*

**S10 Fig.** UMAP representations of selected fibroblast-specific genes across the different stages of wound healing (A) all clusters in unwounded skin (unw) and at 2 dpw, 4 dpw, 6 dpw (B) fibroblast at 4 dpw *fn1b*, *cilp1*, *ogna*, *plaub*, *mmp2, adam8a, hbba1, hbefg*

**S11 Fig.** Violin plots showing the expression levels of ECM build-up and ECM breakdown-related genes in different fibroblast subclusters as well as in all fibroblasts, macrophages and neutrophils together in unwounded skin and across the different phases of cutaneous wound healing.

**S12 Fig.** Contributions of innate immune cells to ECM breakdown UMAP representations of *mmp13a*, *timp2b* (all clusters) in unwounded skin (unw) and at 2 dpw, 4 dpw and 6 dpw

**S13 Fig.** Expression of PDGF and TGFβ1 ligands and receptors in macrophages, neutrophils and fibroblasts UMAP representations (all clusters) of *pdgfab*, *pdgfba*, *pdgfra*, *pdgfrb* and of *tgfb1a*, *tgfb1b*. *tgfbr1a*, *tgfbr1b*, *tgfbr2a*, *tgfbr2b* in unwounded skin (unw) and at 2 dpw, 4 dpw and 8 dpw.

**S14 Fig.** Ligand – receptor analysis between macrophages and fibroblasts. (A) Circos plots showing the secreted signaling pathways from all cell clusters to fibroblasts (upper row) and from macrophages to all cell clusters (lower row) across the time points. (B) Circos plots showing the ECM – receptor signaling pathways from all cell clusters to fibroblasts (upper row) and from macrophages to all cell clusters (lower row) across the time points. (C) Circos plots showing the cell-cell contact signaling pathways from all cell clusters to fibroblasts (upper row) and from macrophages to all cell clusters (lower row) across the time points.

**S15 Fig.** Gain of Lh2 function does not compromise granulation tissue resolution. (A, B) Representative images of immunostaining against collagen I protein (gray) at 8 dpw and 12 dpw on paraffin wound sections of Tg(hsp70l:plod2-p2A-EGFP) fish and their control, counterstained with DAPI (blue). Scale bars: 200 μm.

**S16 Fig.** Genetic loss of Lh2 function does not compromise granulation tissue formation and resolution. (A) H&E staining on sections of *plod2* mutant fish and their control wound sections at 4 dpw, 8 dpw and 12 dpw, respectively (yellow dashed lines mark the granulation tissue). (B) Quantification of the granulation tissue sizes in sections as shown in (A), revealing that sizes were not changed upon loss of *plod2* compared to wild type siblings. Data represent mean ± SD, *n* = 3. (C) Representative images of immunostaining against collagen I protein (gray) at 4 dpw, 8 dpw and 12 dpw on paraffin sections, counterstained with DAPI (blue). (D) Quantification of the collagen I stained areas at 4 dpw, 8 dpw and 12 dpw showing no changes upon loss of *plod2* compared to wild-type siblings. Data represent mean ± SD, *n* = 3. Significances were determined with a two-tailed Student’s t test. (E) TEM analysis of *plod2* mutant fish demonstrated that the diameters and alignment of collagen fibres was not changed at 4 dpw compared to wild-type siblings. Numerical values for panels B and D can be found in S1 Data. Scale bars: 200 μm in A and C; 200 nm in E. ep: epidermis, gt: granulation tissue.

**S1 Data.** Numerical values for panels in Figs 1G, 2B, 3B, 5C,D,H,I,J, 6A, 7A,C,E and S16 Fig B,D.

## References

1. Poss KD, Keating MT, Nechiporuk A. Tales of regeneration in zebrafish. Dev Dyn. 2003;226(2):202–10. doi: 10.1002/dvdy.10220.

2. Larson BJ, Longaker MT, Lorenz HP. Scarless Fetal Wound Healing: A Basic Science Review. Plast Reconstr Surg. 2010;126(4):1172–80. doi: 10.1097/prs.0b013e3181eae781.

3. Lorenz HP. Fetal wound healing. Frontiers in Bioscience. 2003;8(6):s1240–8. doi: 10.2741/1183.

4. Richardson R, Slanchev K, Kraus C, Knyphausen P, Eming S, Hammerschmidt M. Adult Zebrafish as a Model System for Cutaneous Wound-Healing Research. J Invest Dermatol. 2013;133(6):1655–65. doi: 10.1038/jid.2013.16.

5. Tottoli EM, Dorati R, Genta I, Chiesa E, Pisani S, Conti B. Skin Wound Healing Process and New Emerging Technologies for Skin Wound Care and Regeneration. Pharmaceutics. 2020;12(8):735. doi: 10.3390/pharmaceutics12080735.

6. Vannella KM, Wynn TA. Mechanisms of Organ Injury and Repair by Macrophages. Annu Rev Physiol. 2017;79:593–617. doi: 10.1146/annurev-physiol-022516-034356.

7. Willenborg S, Injarabian L, Eming SA. Role of Macrophages in Wound Healing. Cold Spring Harb Perspect Biol. 2022;14(12):a041216. doi: 10.1101/cshperspect.a041216.

8. Wynn TA, Vannella KM. Macrophages in Tissue Repair, Regeneration, and Fibrosis. Immunity. 2016;44:450–62. doi: 10.1016/j.immuni.2016.02.015.

9. Pang J, Maienschein-Cline M, Koh TJ. Monocyte/Macrophage Heterogeneity during Skin Wound Healing in Mice. J Immunol. 2022;209(10):1999–2011. doi: 10.4049/jimmunol.2200365.

10. Willenborg S, Sanin DE, Jais A, Ding X, Ulas T, Nuchel J, et al. Mitochondrial metabolism coordinates stage-specific repair processes in macrophages during wound healing. Cell Metab. 2021;33(12):2398–414 e9. doi: 10.1016/j.cmet.2021.10.004.

11. Krzyszczyk P, Schloss R, Palmer A, Berthiaume F. The Role of Macrophages in Acute and Chronic Wound Healing and Interventions to Promote Pro-wound Healing Phenotypes. Front Physiol. 2018;9. doi: 10.3389/fphys.2018.00419.

12. Schlundt C, Fischer H, Bucher CH, Rendenbach C, Duda GN, Schmidt-Bleek K. The multifaceted roles of macrophages in bone regeneration: A story of polarization, activation and time. Acta Biomater. 2021;133:46–57. doi: 10.1016/j.actbio.2021.04.052.

13. Wang N, Liang H, Zen K. Molecular Mechanisms That Influence the Macrophage M1–M2 Polarization Balance. Front Immunol. 2014;5. doi: 10.3389/fimmu.2014.00614.

14. Fernandes KJL, McKenzie IA, Mill P, Smith KM, Akhavan M, Barnabé-Heider F, et al. A dermal niche for multipotent adult skin-derived precursor cells. Nat Cell Biol. 2004;6(11):1082–93. doi: 10.1038/ncb1181.

15. Plikus MV, Wang X, Sinha S, Forte E, Thompson SM, Herzog EL, et al. Fibroblasts: Origins, definitions, and functions in health and disease. Cell. 2021;184(15):3852–72. doi: 10.1016/j.cell.2021.06.024.

16. Toma JG, Akhavan M, Fernandes KJL, Barnabé-Heider F, Sadikot A, Kaplan DR, et al. Isolation of multipotent adult stem cells from the dermis of mammalian skin. Nat Cell Biol. 2001;3(9):778–84. doi: 10.1038/ncb0901-778.

17. Torregrossa M, Davies L, Hans-Gunther M, Simon JC, Franz S, Rinkevich Y. Effects of embryonic origin, tissue cues and pathological signals on fibroblast diversity in humans. Nat Cell Biol. 2025;27(5):720–35. doi: 10.1038/s41556-025-01638-5.

18. Abe R, Donnelly S, C., Peng T, Bucala R, Metz CN. Peripheral Blood Fibrocytes: Differentiation Pathway and Migration to Wound Sites. J Immunol. 2001;166(12):7556–62. doi: 10.4049/jimmunol.166.12.7556.

19. Pilling D, Gomer RH. Differentiation of circulating monocytes into fibroblast-like cells. Methods Mol Biol. 2012;904:191–206. doi: 10.1007/978-1-61779-943-3_16.

20. Reinhardt JW, Breuer CK. Fibrocytes: A Critical Review and Practical Guide. Front Immunol. 2021;12:784401. doi: 10.3389/fimmu.2021.784401.

21. Reinke JM, Sorg H. Wound Repair and Regeneration. Eur Surg Res. 2012;49(1):35–43. doi: 10.1159/000339613.

22. Hinz B. Formation and Function of the Myofibroblast during Tissue Repair. J Invest Dermatol. 2007;127(3):526–37. doi: 10.1038/sj.jid.5700613.

23. Schuster R, Younesi F, Ezzo M, Hinz B. The Role of Myofibroblasts in Physiological and Pathological Tissue Repair. Cold Spring Harb Perspect Biol. 2023;15(1): a041231. doi: 10.1101/cshperspect.a041231.

24. Correa-Gallegos D, Ye H, Dasgupta B, Sardogan A, Kadri S, Kandi R, et al. CD201(+) fascia progenitors choreograph injury repair. Nature. 2023;623(7988):792–802. doi: 10.1038/s41586-023-06725-x.

25. Jiang D, Guo R, Machens HG, Rinkevich Y. Diversity of Fibroblasts and Their Roles in Wound Healing. Cold Spring Harb Perspect Biol. 2023;15(3). doi: 10.1101/cshperspect.a041222.

26. Jiang D, Rinkevich Y. Distinct fibroblasts in scars and regeneration. Curr Opin Genet Dev. 2021;70:7–14. doi: 10.1016/j.gde.2021.04.005.

27. Mascharak S, desJardins-Park HE, Davitt MF, Griffin M, Borrelli MR, Moore AL, et al. Preventing Engrailed-1 activation in fibroblasts yields wound regeneration without scarring. Science. 2021;372(6540): eaba2374. doi: 10.1126/science.aba2374.

28. Rinkevich Y, Walmsley GG, Hu MS, Maan ZN, Newman AM, Drukker M, et al. Skin fibrosis. Identification and isolation of a dermal lineage with intrinsic fibrogenic potential. Science. 2015;348(6232):aaa2151. doi: 10.1126/science.aaa2151.

29. Sinha S, Sparks HD, Labit E, Robbins HN, Gowing K, Jaffer A, et al. Fibroblast inflammatory priming determines regenerative versus fibrotic skin repair in reindeer. Cell. 2022;185(25):4717–36.e25. doi: 10.1016/j.cell.2022.11.004.

30. Eming SA, Martin P, Tomic-Canic M. Wound repair and regeneration: Mechanisms, signaling, and translation. Sci Transl Med. 2014;6(265):265sr6–sr6. doi: 10.1126/scitranslmed.3009337.

31. Moretti L, Stalfort J, Barker TH, Abebayehu D. The interplay of fibroblasts, the extracellular matrix, and inflammation in scar formation. J Biol Chem. 2022;298(2):101530. doi: 10.1016/j.jbc.2021.101530.

32. Mori R, Shaw TJ, Martin P. Molecular mechanisms linking wound inflammation and fibrosis: knockdown of osteopontin leads to rapid repair and reduced scarring. J Exp Med. 2008;205(1):43–51. doi: 10.1084/jem.20071412.

33. Qi Y, Xu R. Roles of PLODs in Collagen Synthesis and Cancer Progression. Front Cell Dev Biol. 2018;6:66. doi: 10.3389/fcell.2018.00066.

34. Van Der Slot AJ, Zuurmond A-M, Bardoel AFJ, Wijmenga C, Pruijs HEH, Sillence DO, et al. Identification of PLOD2 as Telopeptide Lysyl Hydroxylase, an Important Enzyme in Fibrosis. J Biol Chem. 2003;278(42):40967–72. doi: 10.1074/jbc.m307380200.

35. van der Slot-Verhoeven AJ, van Dura EA, Attema J, Blauw B, Degroot J, Huizinga TW, et al. The type of collagen cross-link determines the reversibility of experimental skin fibrosis. Biochim Biophys Acta. 2005;1740(1):60–7. doi: 10.1016/j.bbadis.2005.02.007.

36. Yamauchi M, Sricholpech M. Lysine post-translational modifications of collagen. Essays Biochem. 2012;52:113–33. doi: 10.1042/bse0520113.

37. Myllyharju J, Kivirikko KI. Collagens, modifying enzymes and their mutations in humans, flies and worms. Trends Genet. 2004;20(1):33–43. doi: 10.1016/j.tig.2003.11.004.

38. Terajima M, Taga Y, Nakamura T, Guo HF, Kayashima Y, Maeda-Smithies N, et al. Lysyl hydroxylase 2 mediated collagen post-translational modifications and functional outcomes. Sci Rep. 2022;12(1):14256. doi: 10.1038/s41598-022-18165-0.

39. Knipper A, Johanna Willenborg S, Brinckmann J, Bloch W, Maaß T, Wagener R, et al. Interleukin-4 Receptor α Signaling in Myeloid Cells Controls Collagen Fibril Assembly in Skin Repair. Immunity. 2015;43(4):803–16. doi: 10.1016/j.immuni.2015.09.005.

40. Richardson R, Metzger M, Knyphausen P, Ramezani T, Slanchev K, Kraus C, et al. Re-epithelialization of cutaneous wounds in adult zebrafish combines mechanisms of wound closure in embryonic and adult mammals. Development. 2016;143(12):2077–88. doi: 10.1242/dev.130492.

41. Raote I, Rosendahl AH, Hakkinen HM, Vibe C, Kucukaylak I, Sawant M, et al. TANGO1 inhibitors reduce collagen secretion and limit tissue scarring. Nat Commun. 2024;15(1):3302. doi: 10.1038/s41467-024-47004-1.

42. Almet AA, Liu Y, Nie Q, Plikus MV. Integrated Single-Cell Analysis Reveals Spatially and Temporally Dynamic Heterogeneity in Fibroblast States during Wound Healing. J Invest Dermatol. 2025;145(3):645–59 e25. doi: 10.1016/j.jid.2024.06.1281.

43. Foster DS, Januszyk M, Yost KE, Chinta MS, Gulati GS, Nguyen AT, et al. Integrated spatial multiomics reveals fibroblast fate during tissue repair. Proc Natl Acad Sci U S A. 2021;118(41). doi: 10.1073/pnas.2110025118.

44. Hu KH, Kuhn NF, Courau T, Tsui J, Samad B, Ha P, et al. Transcriptional space-time mapping identifies concerted immune and stromal cell patterns and gene programs in wound healing and cancer. Cell Stem Cell. 2023;30(6):885–903 e10. doi: 10.1016/j.stem.2023.05.001.

45. Cortada E, Yao J, Xia Y, Dundar F, Zumbo P, Yang B, et al. Cross-species single-cell RNA-seq analysis reveals disparate and conserved cardiac and extracardiac inflammatory responses upon heart injury. Commun Biol. 2024;7(1):1611. doi: 10.1038/s42003-024-07315-x.

46. Denans N, Tran NTT, Swall ME, Diaz DC, Blanck J, Piotrowski T. An anti-inflammatory activation sequence governs macrophage transcriptional dynamics during tissue injury in zebrafish. Nat Commun. 2022;13(1):5356. doi: 10.1038/s41467-022-33015-3.

47. Hu B, Lelek S, Spanjaard B, El-Sammak H, Simões MG, Mintcheva J, et al. Origin and function of activated fibroblast states during zebrafish heart regeneration. Nat Genet. 2022;54(8):1227–37. doi: 10.1038/s41588-022-01129-5.

48. Hou Y, Lee HJ, Chen Y, Ge J, Osman FOI, McAdow AR, et al. Cellular diversity of the regenerating caudal fin. Sci Adv. 2020;6(33):eaba2084. doi: 10.1126/sciadv.aba2084.

49. Wei KH, Lin IT, Chowdhury K, Lim KL, Liu KT, Ko TM, et al. Comparative single-cell profiling reveals distinct cardiac resident macrophages essential for zebrafish heart regeneration. Elife. 2023;12:e84679. doi: 10.7554/eLife.84679.

50. Bevan L, Lim ZW, Venkatesh B, Riley PR, Martin P, Richardson RJ. Specific macrophage populations promote both cardiac scar deposition and subsequent resolution in adult zebrafish. Cardiovasc Res. 2020;116(7):1357–71. doi: 10.1093/cvr/cvz221.

51. Cavone L, Mccann T, Drake LK, Aguzzi EA, Oprişoreanu A-M, Pedersen E, et al. A unique macrophage subpopulation signals directly to progenitor cells to promote regenerative neurogenesis in the zebrafish spinal cord. Dev Cell. 2021;56(11):1617–30.e6. doi: 10.1016/j.devcel.2021.04.031.

52. Hasegawa T, Hall CJ, Crosier PS, Abe G, Kawakami K, Kudo A, et al. Transient inflammatory response mediated by interleukin-1beta is required for proper regeneration in zebrafish fin fold. Elife. 2017;6: e22716. doi: 10.7554/eLife.22716.

53. Jumeau C, Awad F, Assrawi E, Cobret L, Duquesnoy P, Giurgea I, et al. Expression of SAA1, SAA2 and SAA4 genes in human primary monocytes and monocyte-derived macrophages. PLOS ONE. 2019;14(5):e0217005. doi: 10.1371/journal.pone.0217005.

54. Nguyen-Chi M, Laplace-Builhe B, Travnickova J, Luz-Crawford P, Tejedor G, Phan QT, et al. Identification of polarized macrophage subsets in zebrafish. eLife. 2015;4:e07288. doi: 10.7554/elife.07288.

55. Madsen DH, Leonard D, Masedunskas A, Moyer A, Jurgensen HJ, Peters DE, et al. M2-like macrophages are responsible for collagen degradation through a mannose receptor-mediated pathway. J Cell Biol. 2013;202(6):951–66. doi: 10.1083/jcb.201301081.

56. Wang Q, Huang F, Duan X, Cheng H, Zhang C, Li L, et al. The ERbeta-CXCL19/CXCR4-NFkappaB pathway is critical in mediating the E2-induced inflammation response in the orange-spotted grouper (Epinephelus coioides). J Steroid Biochem Mol Biol. 2021;212:105926. doi: 10.1016/j.jsbmb.2021.105926.

57. Wu X, Qian L, Zhao H, Lei W, Liu Y, Xu X, et al. CXCL12/CXCR4: An amazing challenge and opportunity in the fight against fibrosis. Ageing Res Rev. 2023;83:101809. doi: 10.1016/j.arr.2022.101809.

58. Watt FM, Fujiwara H. Cell-Extracellular Matrix Interactions in Normal and Diseased Skin. Cold Spring Harb Perspect Biol. 2011;3(4):a005124-a. doi: 10.1101/cshperspect.a005124.

59. Biggs RM, Bradshaw AD. Organ fibrosis: beyond collagen I expression. Fibroblast phenotype and basement membrane proteins. Am J Physiol Cell Physiol. 2025;328(6):C2023–C2031. doi: 10.1152/ajpcell.00077.2025.

60. Simões FC, Cahill TJ, Kenyon A, Gavriouchkina D, Vieira JM, Sun X, et al. Macrophages directly contribute collagen to scar formation during zebrafish heart regeneration and mouse heart repair. Nat Commun. 2020;11(1):600. doi: 10.1038/s41467-019-14263-2.

61. Mitchell JL, Mutch NJ. Let’s cross-link: diverse functions of the promiscuous cellular transglutaminase factor XIII-A. J Thromb Haemost. 2019;17(1):19–30. doi: 10.1111/jth.14348.

62. Campbell L, Emmerson E, Davies F, Gilliver SC, Krust A, Chambon P, et al. Estrogen promotes cutaneous wound healing via estrogen receptor beta independent of its antiinflammatory activities. J Exp Med. 2010;207(9):1825–33. doi: 10.1084/jem.20100500.

63. Campbell L, Emmerson E, Williams H, Saville CR, Krust A, Chambon P, et al. Estrogen receptor-alpha promotes alternative macrophage activation during cutaneous repair. J Invest Dermatol. 2014;134(9):2447–57. doi: 10.1038/jid.2014.175.

64. Xu S, Xie F, Tian L, Fallah S, Babaei F, Manno SHC, et al. Estrogen accelerates heart regeneration by promoting the inflammatory response in zebrafish. J Endocrinol. 2020;245(1):39–51. doi: 10.1530/JOE-19-0413.

65. Matthews KW, Mueller-Ortiz SL, Wetsel RA. Carboxypeptidase N: a pleiotropic regulator of inflammation. Mol Immunol. 2004;40(11):785–93. doi: 10.1016/j.molimm.2003.10.002.

66. Walter AS, Volkmer E, Gauglitz G, Bocker W, Saller MM. Systematic review of molecular pathways in burn wound healing. Burns. 2023;49(7):1525–33. doi: 10.1016/j.burns.2023.03.006.

67. Hinz B, Pittet P, Smith-Clerc J, Chaponnier C, Meister JJ. Myofibroblast Development Is Characterized by Specific Cell-Cell Adherens Junctions. Mol Biol Cell. 2004;15:4310–20. doi: 10.1091/mbc.E04-05-0386.

68. Mazzeo L, Ghosh S, Di Cicco E, Isma J, Tavernari D, Samarkina A, et al. ANKRD1 is a mesenchymal-specific driver of cancer-associated fibroblast activation bridging androgen receptor loss to AP-1 activation. Nat Commun. 2024;15(1):1038. doi: 10.1038/s41467-024-45308-w.

69. Samaras SE, Almodovar-Garcia K, Wu N, Yu F, Davidson JM. Global deletion of Ankrd1 results in a wound-healing phenotype associated with dermal fibroblast dysfunction. Am J Pathol. 2015;185(1):96–109. doi: 10.1016/j.ajpath.2014.09.018.

70. Kanisicak O, Khalil H, Ivey MJ, Karch J, Maliken BD, Correll RN, et al. Genetic lineage tracing defines myofibroblast origin and function in the injured heart. Nat Commun. 2016;7:12260. doi: 10.1038/ncomms12260.

71. Ricard-Blum S. The collagen family. Cold Spring Harb Perspect Biol. 2011;3(1):a004978. Epub 20110101. doi: 10.1101/cshperspect.a004978.

72. Wehner D, Tsarouchas TM, Michael A, Haase C, Weidinger G, Reimer MM, et al. Wnt signaling controls pro-regenerative Collagen XII in functional spinal cord regeneration in zebrafish. Nat Commun. 2017;8(1):126. doi: 10.1038/s41467-017-00143-0.

73. Lai CS, Tu CW, Kuo HC, Sun PP, Tsai ML. Type II Collagen from Cartilage of Acipenser baerii Promotes Wound Healing in Human Dermal Fibroblasts and in Mouse Skin. Mar Drugs. 2020;18(10):511. doi: 10.3390/md18100511.

74. Baker CE, Moore-Lotridge SN, Hysong AA, Posey SL, Robinette JP, Blum DM, et al. Bone Fracture Acute Phase Response-A Unifying Theory of Fracture Repair: Clinical and Scientific Implications. Clin Rev Bone Miner Metab. 2018;16(4):142–58. doi: 10.1007/s12018-018-9256-x.

75. Han T, Zhu T, Lu Y, Wang Q, Bian H, Chen J, et al. Collagen type X expression and chondrocyte hypertrophic differentiation during OA and OS development. Am J Cancer Res. 2024;14(4):1784–801. doi: 10.62347/JWGW7377.

76. Raman R, Antony M, Nivelle R, Lavergne A, Zappia J, Guerrero-Limon G, et al. The Osteoblast Transcriptome in Developing Zebrafish Reveals Key Roles for Extracellular Matrix Proteins Col10a1a and Fbln1 in Skeletal Development and Homeostasis. Biomolecules. 2024;14(2):139. doi: 10.3390/biom14020139.

77. Leitinger B, Kwan AP. The discoidin domain receptor DDR2 is a receptor for type X collagen. Matrix Biol. 2006;25(6):355–64. doi: 10.1016/j.matbio.2006.05.006.

78. Olaso E, Lin HC, Wang LH, Friedman SL. Impaired dermal wound healing in discoidin domain receptor 2-deficient mice associated with defective extracellular matrix remodeling. Fibrogenesis Tissue Repair. 2011;4(1):5. doi: 10.1186/1755-1536-4-5.

79. Heljasvaara R, Aikio M, Ruotsalainen H, Pihlajaniemi T. Collagen XVIII in tissue homeostasis and dysregulation - Lessons learned from model organisms and human patients. Matrix Biol. 2017;57–58:55-75. doi: 10.1016/j.matbio.2016.10.002.

80. Seppinen L, Sormunen R, Soini Y, Elamaa H, Heljasvaara R, Pihlajaniemi T. Lack of collagen XVIII accelerates cutaneous wound healing, while overexpression of its endostatin domain leads to delayed healing. Matrix Biol. 2008;27(6):535–46. doi: 10.1016/j.matbio.2008.03.003.

81. Dalton CJ, Lemmon CA. Fibronectin: Molecular Structure, Fibrillar Structure and Mechanochemical Signaling. Cells. 2021;10(9):2443. doi: 10.3390/cells10092443.

82. Zhang CL, Zhao Q, Liang H, Qiao X, Wang JY, Wu D, et al. Cartilage intermediate layer protein-1 alleviates pressure overload-induced cardiac fibrosis via interfering TGF-beta1 signaling. J Mol Cell Cardiol. 2018;116:135–44. doi: 10.1016/j.yjmcc.2018.02.006.

83. Nulali J, Zhan M, Zhang K, Tu P, Liu Y, Song H. Osteoglycin: An ECM Factor Regulating Fibrosis and Tumorigenesis. Biomolecules. 2022;12(11):1674. doi: 10.3390/biom12111674.

84. Zou ML, Teng YY, Chen ZH, Liu SY, Jia Y, Zhang KW, et al. The uPA System Differentially Alters Fibroblast Fate and Profibrotic Ability in Skin Fibrosis. Front Immunol. 2022;13:845956. doi: 10.3389/fimmu.2022.845956.

85. Rohani MG, Parks WC. Matrix remodeling by MMPs during wound repair. Matrix Biol. 2015;44–46:113-21. doi: 10.1016/j.matbio.2015.03.002.

86. Mierke CT. The versatile roles of ADAM8 in cancer cell migration, mechanics, and extracellular matrix remodeling. Front Cell Dev Biol. 2023;11:1130823. doi: 10.3389/fcell.2023.1130823.

87. Loo AEK, Wong YT, Ho R, Wasser M, Du T, Ng WT, et al. Effects of Hydrogen Peroxide on Wound Healing in Mice in Relation to Oxidative Damage. PLoS ONE. 2012;7(11):e49215. doi: 10.1371/journal.pone.0049215.

88. Ito J-i, Nagayasu Y, Hoshikawa M, kato KH, Miura Y, Asai K, et al. Enhancement of FGF-1 release along with cytosolic proteins from rat astrocytes by hydrogen peroxide. Brain Res. 2013;1522:12–21. doi: 10.1016/j.brainres.2013.05.035.

89. Singhal SS, Singh SP, Singhal P, Horne D, Singhal J, Awasthi S. Antioxidant role of glutathione S-transferases: 4-Hydroxynonenal, a key molecule in stress-mediated signaling. Toxicol Appl Pharmacol. 2015;289(3):361–70. doi: 10.1016/j.taap.2015.10.006.

90. Widmer CC, Pereira CP, Gehrig P, Vallelian F, Schoedon G, Buehler PW, et al. Hemoglobin can attenuate hydrogen peroxide-induced oxidative stress by acting as an antioxidative peroxidase. Antioxid Redox Signal. 2010;12(2):185–98. doi: 10.1089/ars.2009.2826.

91. Petrov PB, Considine JM, Izzi V, Naba A. Matrisome AnalyzeR – a suite of tools to annotate and quantify ECM molecules in big datasets across organisms. J Cell Sci. 2023;136(17):jcs261255. doi: 10.1242/jcs.261255.

92. Wyatt RA, Keow JY, Harris ND, Hache CA, Li DH, Crawford BD. The zebrafish embryo: a powerful model system for investigating matrix remodeling. Zebrafish. 2009;6(4):347–54. doi: 10.1089/zeb.2009.0609.

93. Maskos K, Lang R, Tschesche H, Bode W. Flexibility and variability of TIMP binding: X-ray structure of the complex between collagenase-3/MMP-13 and TIMP-2. J Mol Biol. 2007;366(4):1222–31. doi: 10.1016/j.jmb.2006.11.072.

94. Singer AJ, Clark RA. Cutaneous wound healing. N Engl J Med. 1999;341(10):738–46. doi: 10.1056/NEJM199909023411006.

95. Bonner JC. Regulation of PDGF and its receptors in fibrotic diseases. Cytokine Growth Factor Rev. 2004;15:255–73. doi: 10.1016/j.cytogfr.2004.03.006.

96. Bauer J, S N, Reisinger F, Zöller J, Yuan D, Heikenwälder M. Lymphotoxin, NF-ĸB, and cancer: the dark side of cytokines. Dig Dis. 2012;30:453–68. doi: 10.1159/000341690.

97. Steele H, Cheng J, Willicut A, Dell G, Breckenridge J, Culberson E, et al. TNF superfamily control of tissue remodeling and fibrosis. Front Immunol 2023;14:1219907. doi: 10.3389/fimmu.2023.1219907.

98. Hardman MJ, Waite A, Zeef L, Burow M, Nakayama T, Ashcroft GS. Macrophage migration inhibitory factor: a central regulator of wound healing. Am J Pathol. 2005;167(6):1561–74. doi: 10.1016/S0002-9440(10)61241-2.

99. Zhu Y, Chen L, Song B, Cui Z, Chen G, Yu Z, et al. Insulin-like Growth Factor-2 (IGF-2) in Fibrosis. Biomolecules. 2022;12:1557. doi: 10.3390/biom12111557.

100. Jian J, Konopka J, Liu C. Insights into the role of progranulin in immunity, infection, and inflammation. J Leukoc Biol. 2013;93:199–208. doi: 10.1189/jlb.0812429.

101. Kerrigan AM, Dennehy KM, Mourão-Sá D, Faro-Trindade I, Willment JA, Taylor PR, et al. CLEC-2 is a phagocytic activation receptor expressed on murine peripheral blood neutrophils. J Immunol. 2009;182:4150–7. doi: 10.4049/jimmunol.0802808.

102. Klingensmith NJ, Fay KT, Swift DA, Bazzano JM, Lyons JD, Chen CW, et al. Junctional adhesion molecule-A deletion increases phagocytosis and improves survival in a murine model of sepsis. JCI Insight. 2022;7:e156255. doi: 10.1172/jci.insight.156255.

103. Hu Q, Tan H, Irwin DM. Evolution of the Vertebrate Resistin Gene Family. PLoS One. 2015;10(6):e0130188. doi: 10.1371/journal.pone.0130188.

104. Schneider VA, Granato M. Genomic structure and embryonic expression of zebrafish lysyl hydroxylase 1 and lysyl hydroxylase 2. Matrix Biol. 2007;26(1):12–9. doi: 10.1016/j.matbio.2006.09.007.

105. Gjaltema RAF, De Rond S, Rots MG, Bank RA. Procollagen Lysyl Hydroxylase 2 Expression Is Regulated by an Alternative Downstream Transforming Growth Factor β-1 Activation Mechanism. J Biol Chem. 2015;290(47):28465–76. doi: 10.1074/jbc.m114.634311.

106. Xu F, Zhang J, Hu G, Liu L, Liang W. Hypoxia and TGF-β1 induced PLOD2 expression improve the migration and invasion of cervical cancer cells by promoting epithelial-to-mesenchymal transition (EMT) and focal adhesion formation. Cancer Cell Int. 2017;17(1). doi: 10.1186/s12935-017-0420-z.

107. Rosell-García T, Palomo-Álvarez O, Rodríguez-Pascual F. A hierarchical network of hypoxia-inducible factor and SMAD proteins governs procollagen lysyl hydroxylase 2 induction by hypoxia and transforming growth factor β1. J Biol Chem. 2019;294(39):14308–18. doi: 10.1074/jbc.ra119.007674.

108. Wen G, Hong M, Li B, Liao W, Cheng SK, Hu B, et al. Transforming growth factor-β-induced protein (TGFBI) suppresses mesothelioma progression through the Akt/mTOR pathway. Int J Oncol. 2011;39(4):1001–9. doi: 10.3892/ijo.2011.1097.

109. Yue W, Zhang H, Gao Y, Ding J, Xue R, Dong C, et al. Procollagen-lysine 2-oxoglutarate 5-dioxygenase 2 promotes collagen cross-linking and ECM stiffening to induce liver fibrosis. Biochim Biophys Acta Mol Basis Dis. 2024;1870(5):167205. doi: 10.1016/j.bbadis.2024.167205.

110. Gistelinck C, Witten PE, Huysseune A, Symoens S, Malfait F, Larionova D, et al. Loss of Type I Collagen Telopeptide Lysyl Hydroxylation Causes Musculoskeletal Abnormalities in a Zebrafish Model of Bruck Syndrome. J Bone Miner Res. 2016;31(11):1930–42. doi: 10.1002/jbmr.2977.

111. Gonzalez-Rosa JM, Martin V, Peralta M, Torres M, Mercader N. Extensive scar formation and regression during heart regeneration after cryoinjury in zebrafish. Development. 2011;138(9):1663–74. doi: 10.1242/dev.060897.

112. Sorrell JM, Caplan AI. Fibroblast heterogeneity: more than skin deep. J Cell Sci. 2004;117(5):667–75. doi: 10.1242/jcs.01005.

113. Wang X, Copmans D, de Witte PAM. Using Zebrafish as a Disease Model to Study Fibrotic Disease. Int J Mol Sci. 2021;22(12):6404. doi: 10.3390/ijms22126404.

114. Ibrahim MM, Chen L, Bond JE, Medina MA, Ren L, Kokosis G, et al. Myofibroblasts contribute to but are not necessary for wound contraction. Lab Invest. 2015;95:1429–38. doi: 10.1038/labinvest.2015.116.

115. Chablais F, Veit J, Rainer G, Jazwinska A. The zebrafish heart regenerates after cryoinjury-induced myocardial infarction. BMC Dev Biol. 2011;11:21. doi: 10.1186/1471-213X-11-21.

116. Gonzalez-Rosa JM, Peralta M, Mercader N. Pan-epicardial lineage tracing reveals that epicardium derived cells give rise to myofibroblasts and perivascular cells during zebrafish heart regeneration. Dev Biol. 2012;370(2):173–86. doi: 10.1016/j.ydbio.2012.07.007.

117. Wang R, Zhang P, Wang J, Ma L, E W, Suo S, et al. Construction of a cross-species cell landscape at single-cell level. Nucleic Acids Res. 2023;51(2):501–16. doi: 10.1093/nar/gkac633.

118. Sánchez-Iranzo H, Galardi-Castilla M, Sanz-Morejón A, González-Rosa JM, Costa R, Ernst A, et al. Transient fibrosis resolves via fibroblast inactivation in the regenerating zebrafish heart. Proc Natl Acad Sci U S A. 2018;115(16):4188–93. doi: 10.1073/pnas.1716713115.

119. Kettleborough RNW, Busch-Nentwich EM, Harvey SA, Dooley CM, De Bruijn E, Van Eeden F, et al. A systematic genome-wide analysis of zebrafish protein-coding gene function. Nature. 2013;496(7446):494–7. doi: 10.1038/nature11992.

120. Löhr H, Hess S, Pereira MMA, Reinoß P, Leibold S, Schenkel C, et al. Diet-Induced Growth Is Regulated via Acquired Leptin Resistance and Engages a Pomc-Somatostatin-Growth Hormone Circuit. Cell Rep. 2018;23(6):1728–41. doi: 10.1016/j.celrep.2018.04.018.

121. Brinckmann J, Kim S, Wu J, Reinhardt DP, Batmunkh C, Metzen E, et al. Interleukin 4 and prolonged hypoxia induce a higher gene expression of lysyl hydroxylase 2 and an altered cross-link pattern: important pathogenetic steps in early and late stage of systemic scleroderma? Matrix Biol. 2005;24(7):459–68. doi: 10.1016/j.matbio.2005.07.002.

122. Gharbia FZ, Abouhashem AS, Moqidem YA, Elbaz AA, Abdellatif A, Singh K, et al. Adult skin fibroblast state change in murine wound healing. Sci Rep. 2023;13(1):886. doi: 10.1038/s41598-022-27152-4.

123. He Y, Kim J, Tacconi C, Moody J, Dieterich LC, Anzengruber F, et al. Mediators of Capillary-to-Venule Conversion in the Chronic Inflammatory Skin Disease Psoriasis. J Invest Dermatol. 2022;142(12):3313–26 e13. doi: 10.1016/j.jid.2022.05.1089.

124. Liu M, Liu X, Zhang J, Liang S, Gong Y, Shi S, et al. Single-cell RNA sequencing reveals the heterogeneity of myofibroblasts in wound repair. Genomics. 2025;117(1):110982. doi: 10.1016/j.ygeno.2024.110982.

125. Shim J, Oh SJ, Yeo E, Park JH, Bae JH, Kim SH, et al. Integrated Analysis of Single-Cell and Spatial Transcriptomics in Keloids: Highlights on Fibrovascular Interactions in Keloid Pathogenesis. J Invest Dermatol. 2022;142(8):2128–39 e11. doi: 10.1016/j.jid.2022.01.017.

126. Yu G, Wang LG, Han Y, He QY. clusterProfiler: an R package for comparing biological themes among gene clusters. OMICS. 2012;16(5):284–7. Epub 20120328. doi: 10.1089/omi.2011.0118.

127. Jin S, Guerrero-Juarez CF, Zhang L, Chang I, Ramos R, Kuan CH, et al. Inference and analysis of cell-cell communication using CellChat. Nat Commun. 2021;12(1):1088. doi: 10.1038/s41467-021-21246-9. P

128. Szklarczyk D, Kirsch R, Koutrouli M, Nastou K, Mehryary F, Hachilif R, et al. The STRING database in 2023: protein–protein association networks and functional enrichment analyses for any sequenced genome of interest. Nucleic Acids Res. 2023;51(D1):D638–D46. doi: 10.1093/nar/gkac1000.

