## Supporting Figures for "Exploring mechanisms of scar-free skin wound healing in adult zebrafish in comparison to mouse"

S1 Fig.

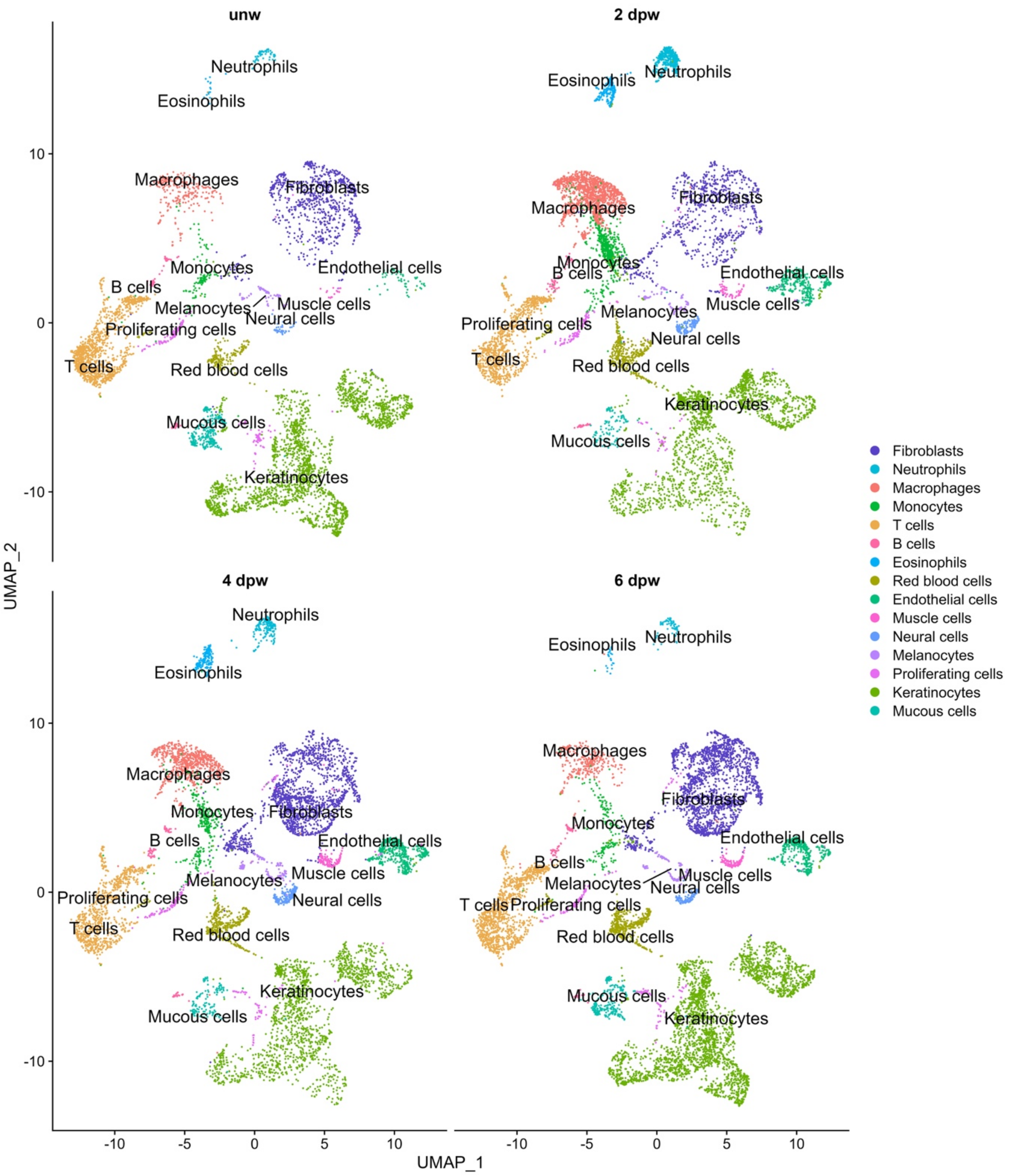
unw
2 dpw
4 dpw
6 dpw
Neutrophils
Eosinophils
Macrophages
Fibroblasts
Monocytes
Endothelial cells
B cells
Melanocytes
Muscle cells
Neural cells
Proliferating cells
T cells
Red blood cells
Mucous cells
Keratinocytes
10
0
-10
-10
-5
0
5
10
UMAP_1
UMAP_2
Fibroblasts
Neutrophils
Macrophages
Monocytes
T cells
B cells
Eosinophils
Red blood cells
Endothelial cells
Muscle cells
Neural cells
Melanocytes
Proliferating cells
Keratinocytes
Mucous cells

S2 Fig.

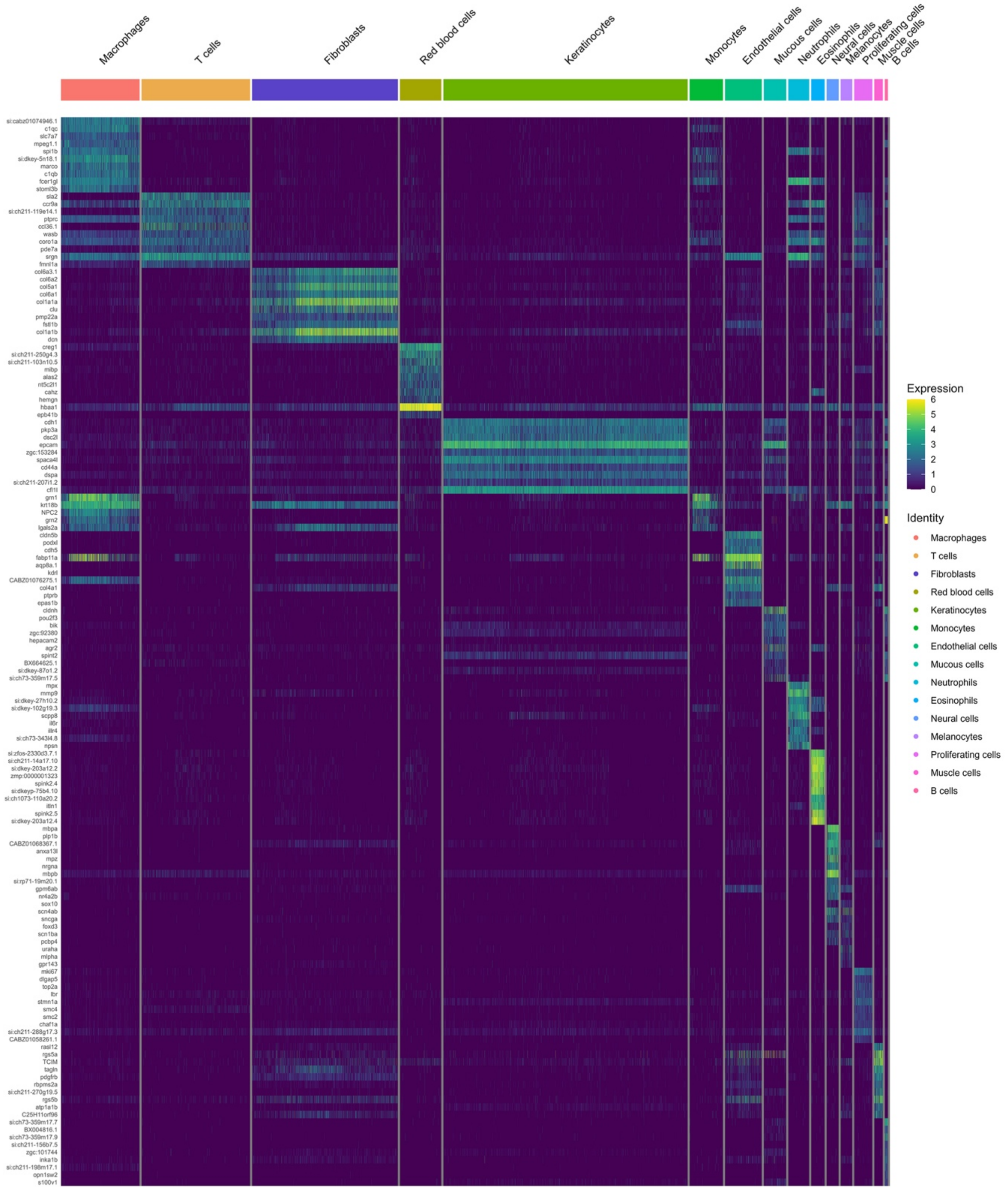
Macrophages
T cells
Fibroblasts
Red blood cells
Keratinocytes
Monocytes
Endothelial cells
Mucous cells
Neutrophils
Eosinophils
Neural cells
Melanocytes
Proliferating cells
Muscle cells
B cells
Expression
6
5
4
3
2
1
0
Identity
Macrophages
T cells
Fibroblasts
Red blood cells
Keratinocytes
Monocytes
Endothelial cells
Mucous cells
Neutrophils
Eosinophils
Neural cells
Melanocytes
Proliferating cells
Muscle cells
B cells

### S3 Fig.

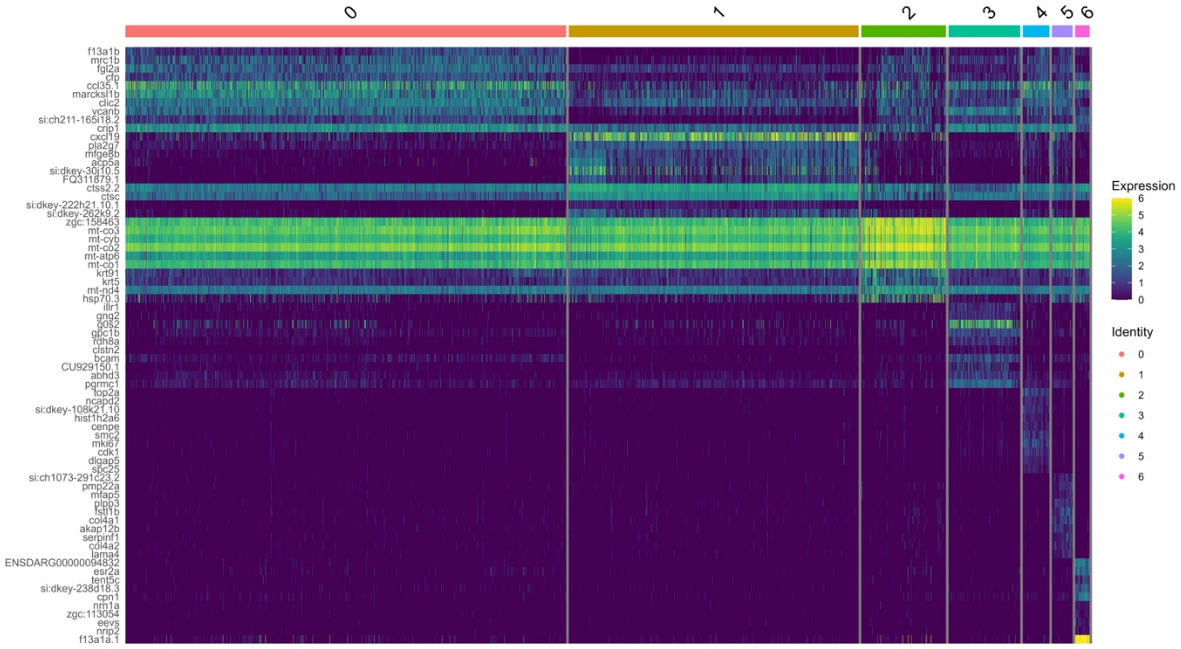
0
1
2
3
4
5
6
f13a1b
mrc1b
fgl2a
cfp
ccl35.1
marcksl1b
clic2
vcanb
si:ch211-165i18.2
crip1
cxcl19
pla2g7
mfge8b
acp5a
si:dkey-30j10.5
FQ311879.1
ctss2.2
ctsc
si:dkey-222h21.10.1
si:dkey-262k9.2
zgc:158463
mt-co3
mt-cyb
mt-co2
mt-atp6
mt-co1
krt91
krt5
mt-nd4
hsp70.3
illr1
gng2
g0s2
gpc1b
rdh8a
clstn2
bcam
CU929150.1
abhd3
pgrmc1
top2a
ncapd2
si:dkey-108k21.10
hist1h2a6
cenpe
smc2
mki67
cdk1
dlgap5
spc25
si:ch1073-291c23.2
pmp22a
mfap5
plpp3
fstl1b
col4a1
akap12b
serpinf1
col4a2
lama4
ENSDARG00000094832
esr2a
tent5c
si:dkey-238d18.3
cpn1
nrn1a
zgc:113054
eevs
nrip2
f13a1a.1
Expression
6
5
4
3
2
1
0
Identity
0
1
2
3
4
5
6

S4 Fig.

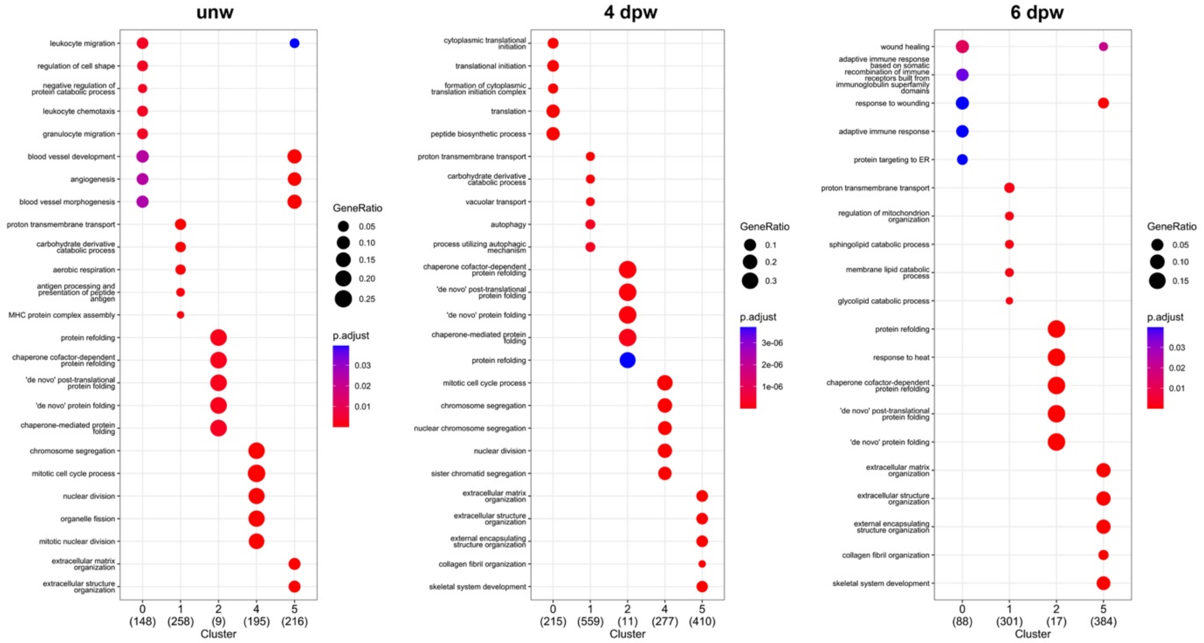
unw
leukocyte migration
regulation of cell shape
negative regulation of protein catabolic process
leukocyte chemotaxis
granulocyte migration
blood vessel development
angiogenesis
blood vessel morphogenesis
proton transmembrane transport
carbohydrate derivative catabolic process
aerobic respiration
antigen processing and presentation of peptide antigen
MHC protein complex assembly
protein refolding
chaperone cofactor-dependent protein refolding
'de novo' post-translational protein folding
'de novo' protein folding
chaperone-mediated protein folding
chromosome segregation
mitotic cell cycle process
nuclear division
organelle fission
mitotic nuclear division
extracellular matrix organization
extracellular structure organization
0 (148)
1 (258)
2 (9)
4 (195)
5 (216)
Cluster
GeneRatio
0.05
0.10
0.15
0.20
0.25
p.adjust
0.03
0.02
0.01
4 dpw
cytoplasmic translational initiation
translational initiation
formation of cytoplasmic translation initiation complex
translation
peptide biosynthetic process
proton transmembrane transport
carbohydrate derivative catabolic process
vacuolar transport
autophagy
process utilizing autophagic mechanism
chaperone cofactor-dependent protein refolding
'de novo' post-translational protein folding
'de novo' protein folding
chaperone-mediated protein folding
protein refolding
mitotic cell cycle process
chromosome segregation
nuclear chromosome segregation
nuclear division
sister chromatid segregation
extracellular matrix organization
extracellular structure organization
external encapsulating structure organization
collagen fibril organization
skeletal system development
0 (215)
1 (559)
2 (11)
4 (277)
5 (410)
Cluster
GeneRatio
0.1
0.2
0.3
p.adjust
3e-06
2e-06
1e-06
6 dpw
wound healing
adaptive immune response based on somatic recombination of immune receptors built from immunoglobulin superfamily domains
response to wounding
adaptive immune response
protein targeting to ER
proton transmembrane transport
regulation of mitochondrion organization
sphingolipid catabolic process
membrane lipid catabolic process
glycolipid catabolic process
protein refolding
response to heat
chaperone cofactor-dependent protein refolding
'de novo' post-translational protein folding
'de novo' protein folding
extracellular matrix organization
extracellular structure organization
external encapsulating structure organization
collagen fibril organization
skeletal system development
0 (88)
1 (301)
2 (17)
5 (384)
Cluster
GeneRatio
0.05
0.10
0.15
p.adjust
0.03
0.02
0.01

S5 Fig.

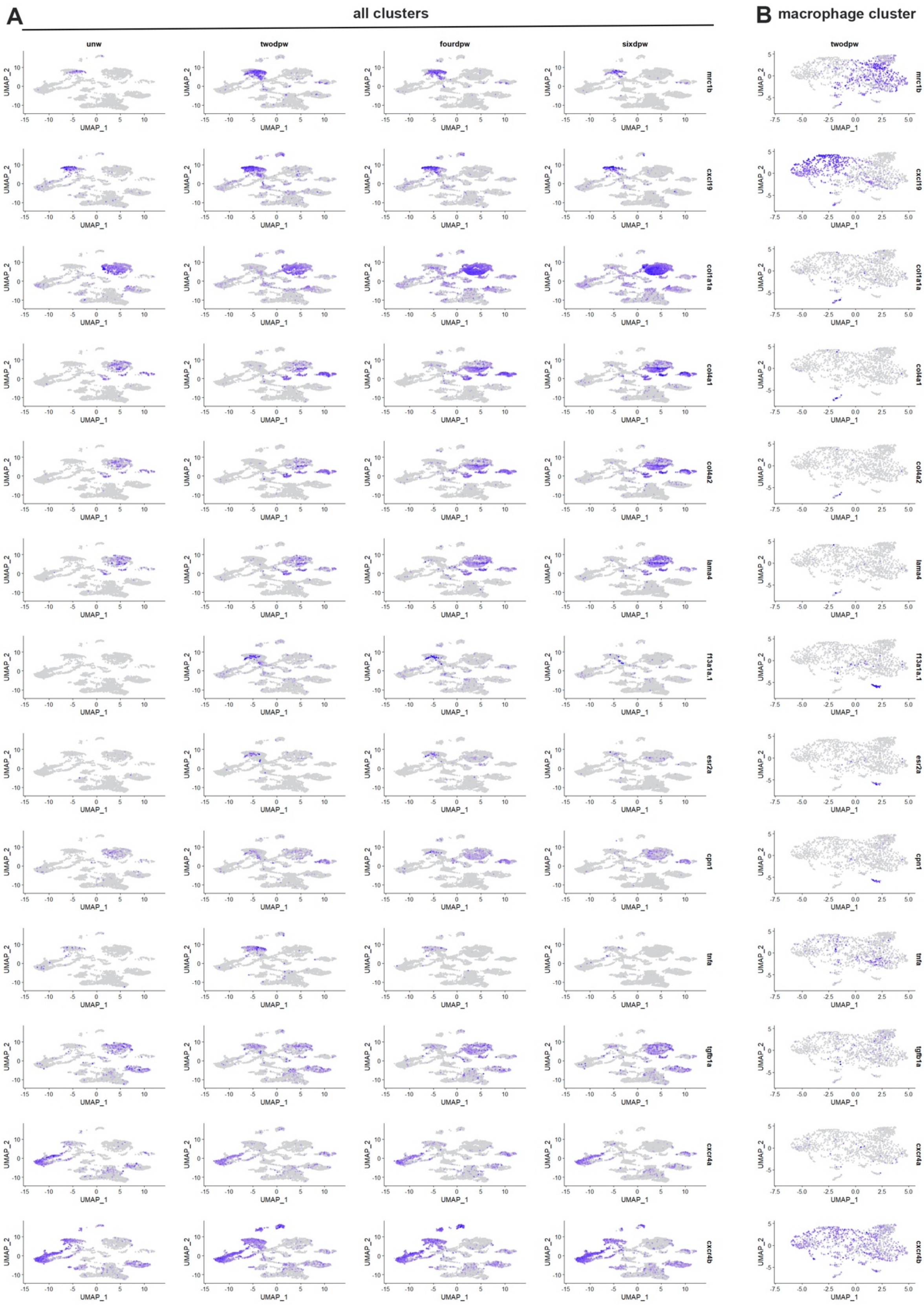

A
all clusters
unw
twodpw
fourdpw
sixdpw
B
macrophage cluster
twodpw
mrc1b
cxcl19
col1a1a
col4a1
col4a2
lama4
f13a1a.1
esr2a
cpn1
tnfa
tgfb1a
cxcr4a
cxcr4b
UMAP_1
UMAP_2

S6 Fig.

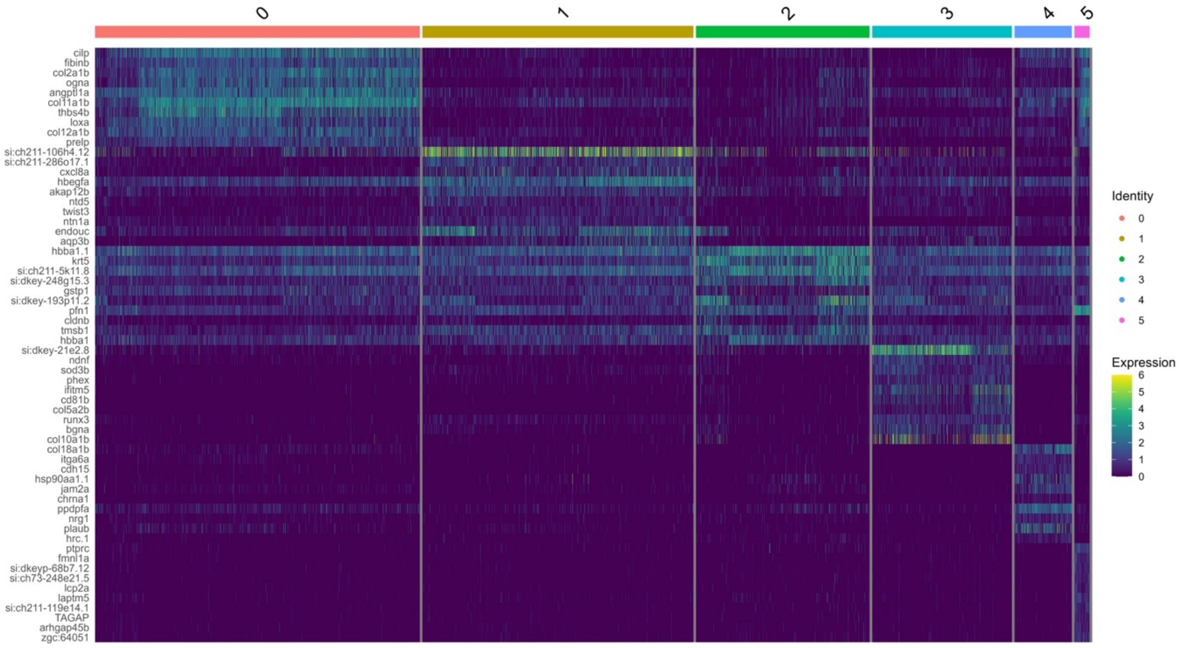
0
1
2
3
4
5
cilp
fibinb
col2a1b
ogna
angptl1a
col11a1b
thbs4b
loxa
col12a1b
prelp
si:ch211-106h4.12
si:ch211-286o17.1
cxcl8a
hbegfa
akap12b
ntd5
twist3
ntn1a
endouc
aqp3b
hbba1.1
krt5
si:ch211-5k11.8
si:dkey-248g15.3
gstp1
si:dkey-193p11.2
pfn1
cldnb
tmsb1
hbba1
si:dkey-21e2.8
ndnf
sod3b
phex
ifitm5
cd81b
col5a2b
runx3
bgna
col10a1b
col18a1b
itga6a
cdh15
hsp90aa1.1
jam2a
chrna1
ppdpfa
nrg1
plaub
hrc.1
ptprc
fmnl1a
si:dkeyp-68b7.12
si:ch73-248e21.5
lcp2a
laptm5
si:ch211-119e14.1
TAGAP
arhgap45b
zgc:64051
Identity
0
1
2
3
4
5
Expression
6
5
4
3
2
1
0

S7 Fig.

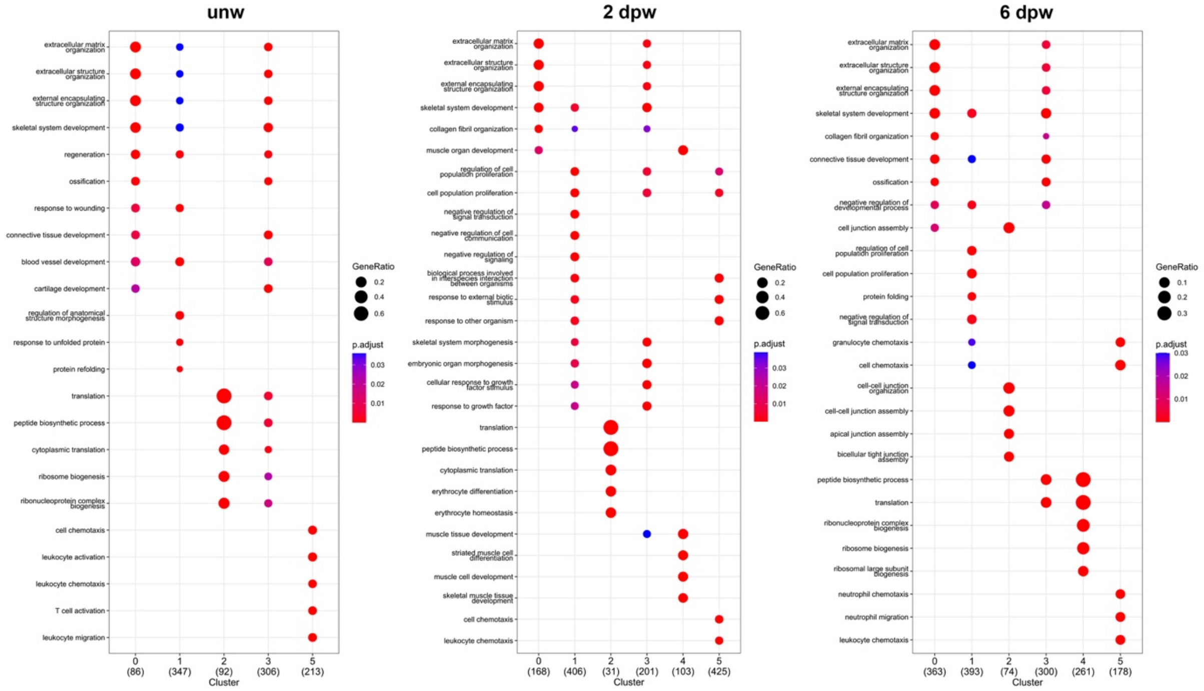
unw
extracellular matrix organization
extracellular structure organization
external encapsulating structure organization
skeletal system development
regeneration
ossification
response to wounding
connective tissue development
blood vessel development
cartilage development
regulation of anatomical structure morphogenesis
response to unfolded protein
protein refolding
translation
peptide biosynthetic process
cytoplasmic translation
ribosome biogenesis
ribonucleoprotein complex biogenesis
cell chemotaxis
leukocyte activation
leukocyte chemotaxis
T cell activation
leukocyte migration
0 (86) 1 (347) 2 (92) 3 (306) 5 (213)
Cluster
GeneRatio
0.2
0.4
0.6
p.adjust
0.03
0.02
0.01
2 dpw
extracellular matrix organization
extracellular structure organization
external encapsulating structure organization
skeletal system development
collagen fibril organization
muscle organ development
regulation of cell population proliferation
cell population proliferation
negative regulation of signal transduction
negative regulation of cell communication
negative regulation of signaling
biological process involved in interspecies interaction between organisms
response to external biotic stimulus
response to other organism
skeletal system morphogenesis
embryonic organ morphogenesis
cellular response to growth factor stimulus
response to growth factor
translation
peptide biosynthetic process
cytoplasmic translation
erythrocyte differentiation
erythrocyte homeostasis
muscle tissue development
striated muscle cell differentiation
muscle cell development
skeletal muscle tissue development
cell chemotaxis
leukocyte chemotaxis
0 (168) 1 (406) 2 (31) 3 (201) 4 (103) 5 (425)
Cluster
GeneRatio
0.2
0.4
0.6
p.adjust
0.03
0.02
0.01
6 dpw
extracellular matrix organization
extracellular structure organization
external encapsulating structure organization
skeletal system development
collagen fibril organization
connective tissue development
ossification
negative regulation of developmental process
cell junction assembly
regulation of cell population proliferation
cell population proliferation
protein folding
negative regulation of signal transduction
granulocyte chemotaxis
cell chemotaxis
cell-cell junction organization
cell-cell junction assembly
apical junction assembly
bicellular tight junction assembly
peptide biosynthetic process
translation
ribonucleoprotein complex biogenesis
ribosome biogenesis
ribosomal large subunit biogenesis
neutrophil chemotaxis
neutrophil migration
leukocyte chemotaxis
0 (363) 1 (393) 2 (74) 3 (300) 4 (261) 5 (178)
Cluster
GeneRatio
0.1
0.2
0.3
p.adjust
0.03
0.02
0.01

.

S8 Fig.

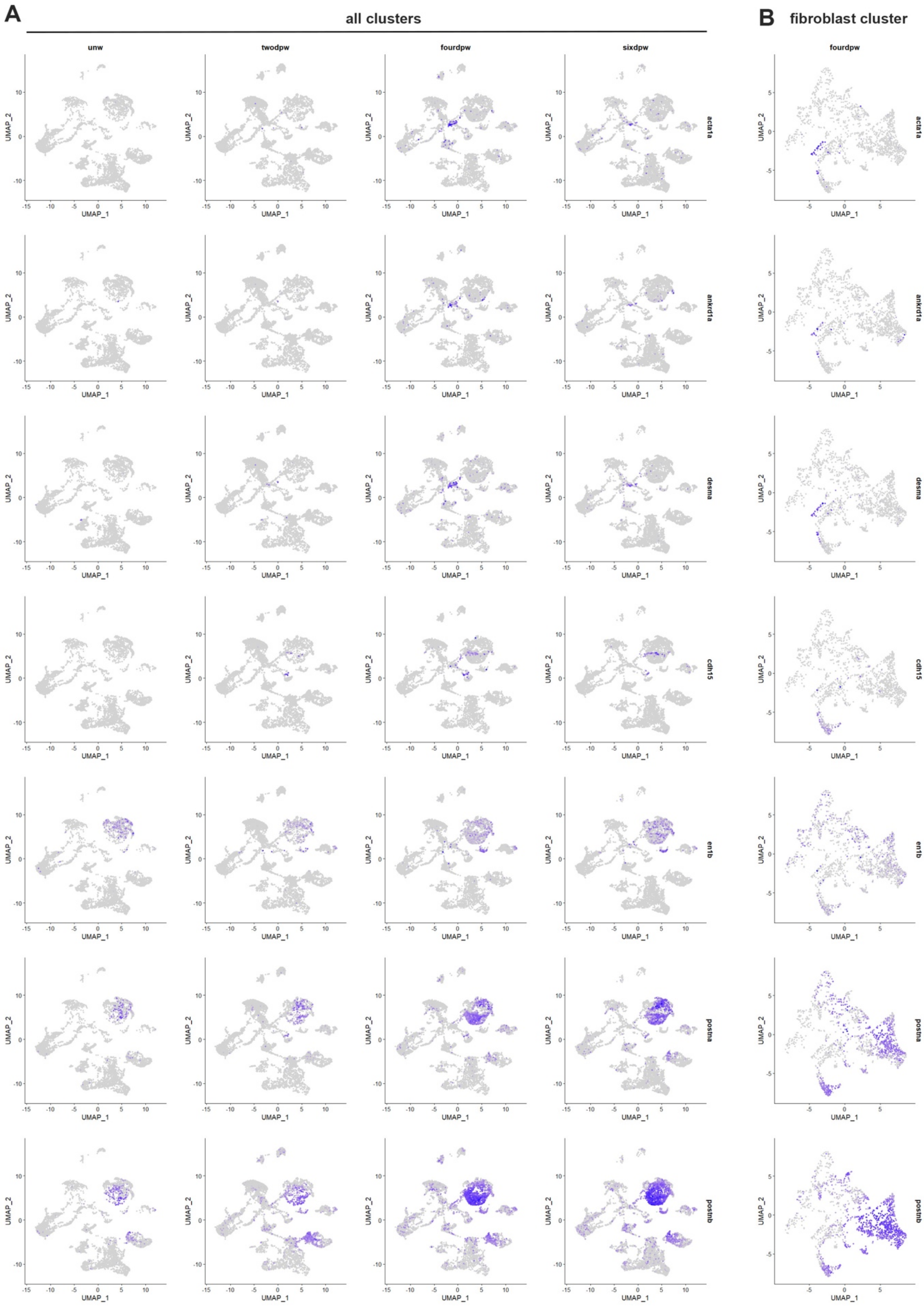
A
all clusters
B
fibroblast cluster
unw
twodpw
fourdpw
sixdpw
fourdpw
acta1a
ankrd1a
desma
cdh15
en1b
postna
postnb
UMAP_1
UMAP_2

S9 Fig.

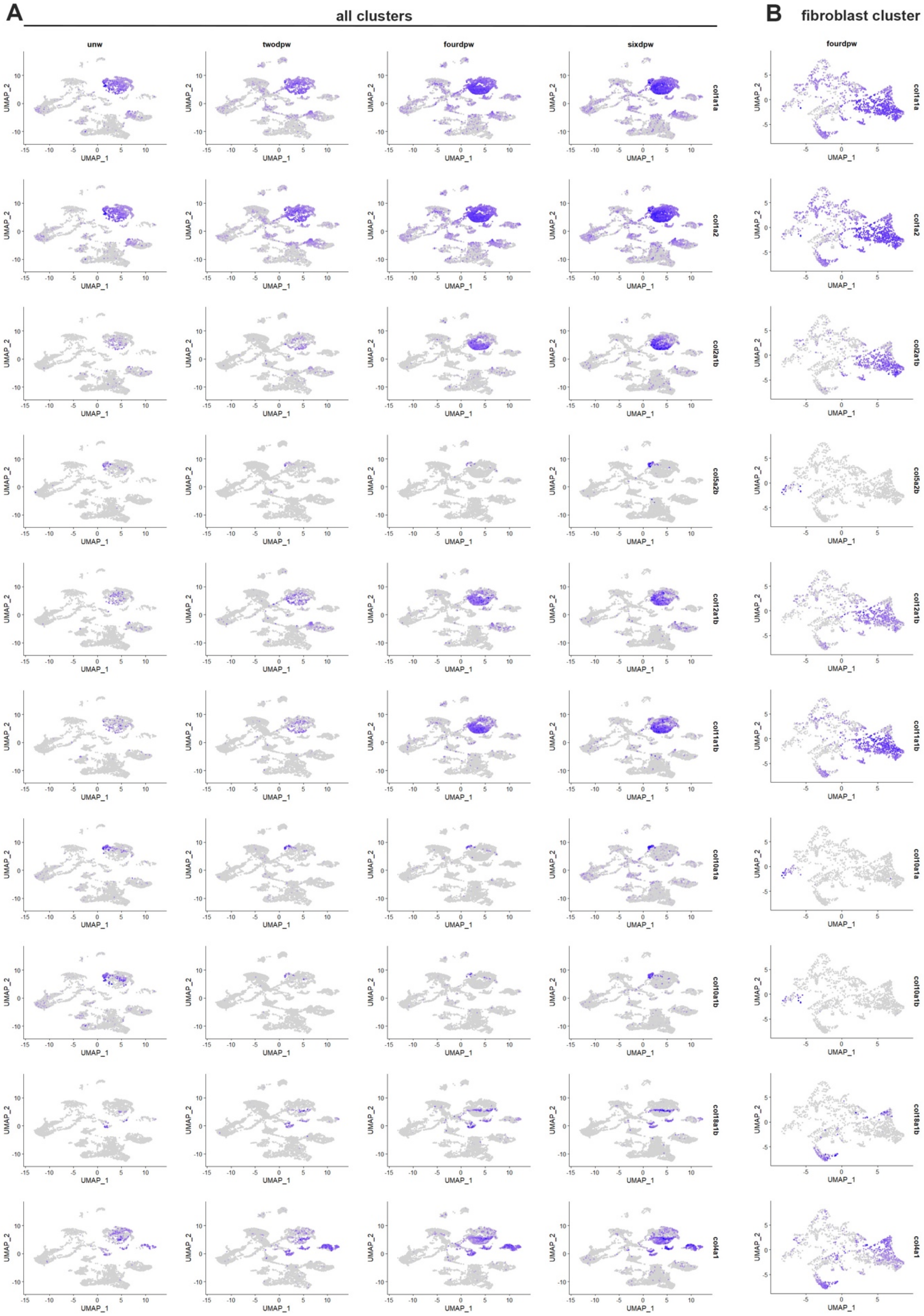
A
all clusters
B
fibroblast cluster
unw
twodpw
fourdpw
sixdpw
fourdpw
col1a1a
col1a2
col2a1b
col5a2b
col12a1b
col11a1b
col10a1a
col10a1b
col8a1b
col4a1
UMAP_1
UMAP_2

S10 Fig.

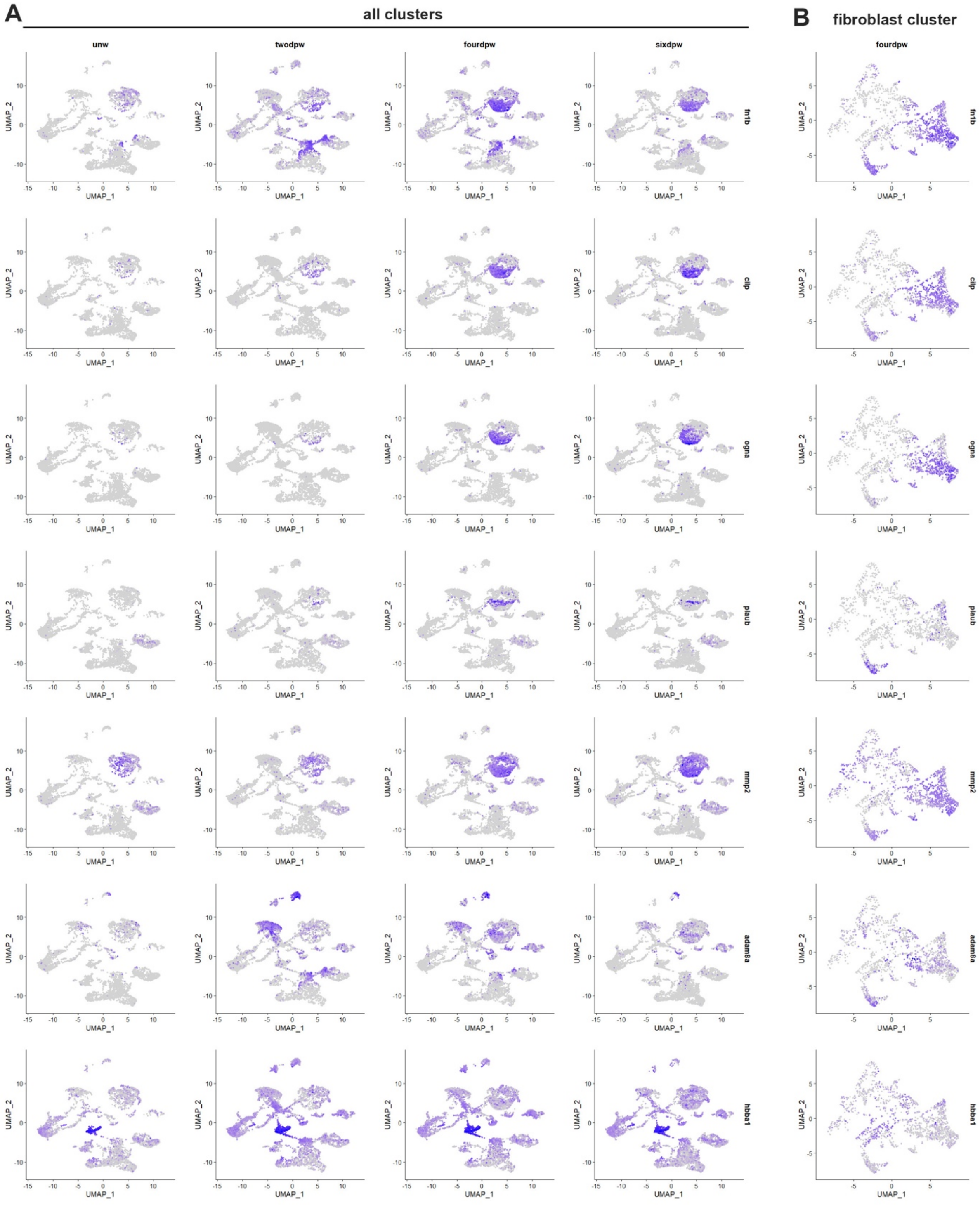
A
all clusters
B
fibroblast cluster
unw
twodpw
fourdpw
sixdpw
fourdpw
fn1b
clip
ogna
plaub
mmp2
adam8a
hbba1
UMAP_1
UMAP_2

S11 Fig.

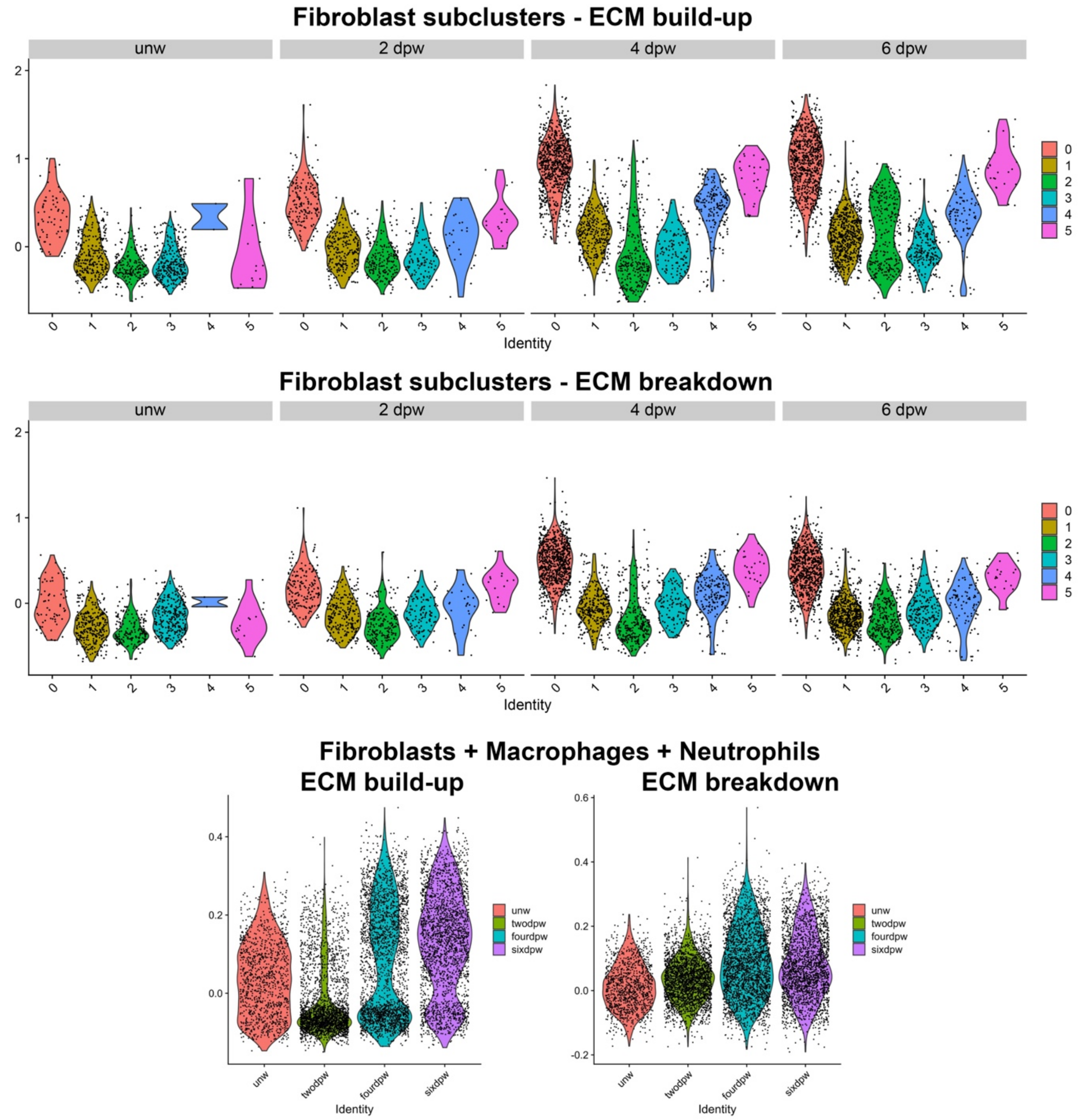
Fibroblast subclusters - ECM build-up
unw
2 dpw
4 dpw
6 dpw
Identity
Fibroblast subclusters - ECM breakdown
Identity
Fibroblasts + Macrophages + Neutrophils
ECM build-up
ECM breakdown
unw
twodpw
fourdpw
sixdpw
Identity

S12 Fig.

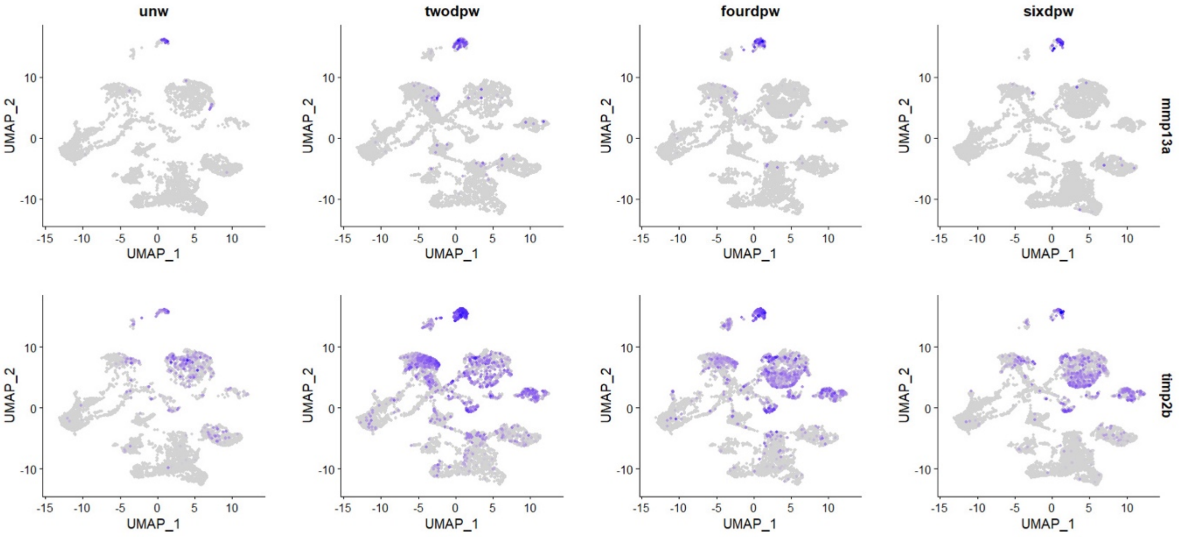

S13 Fig.

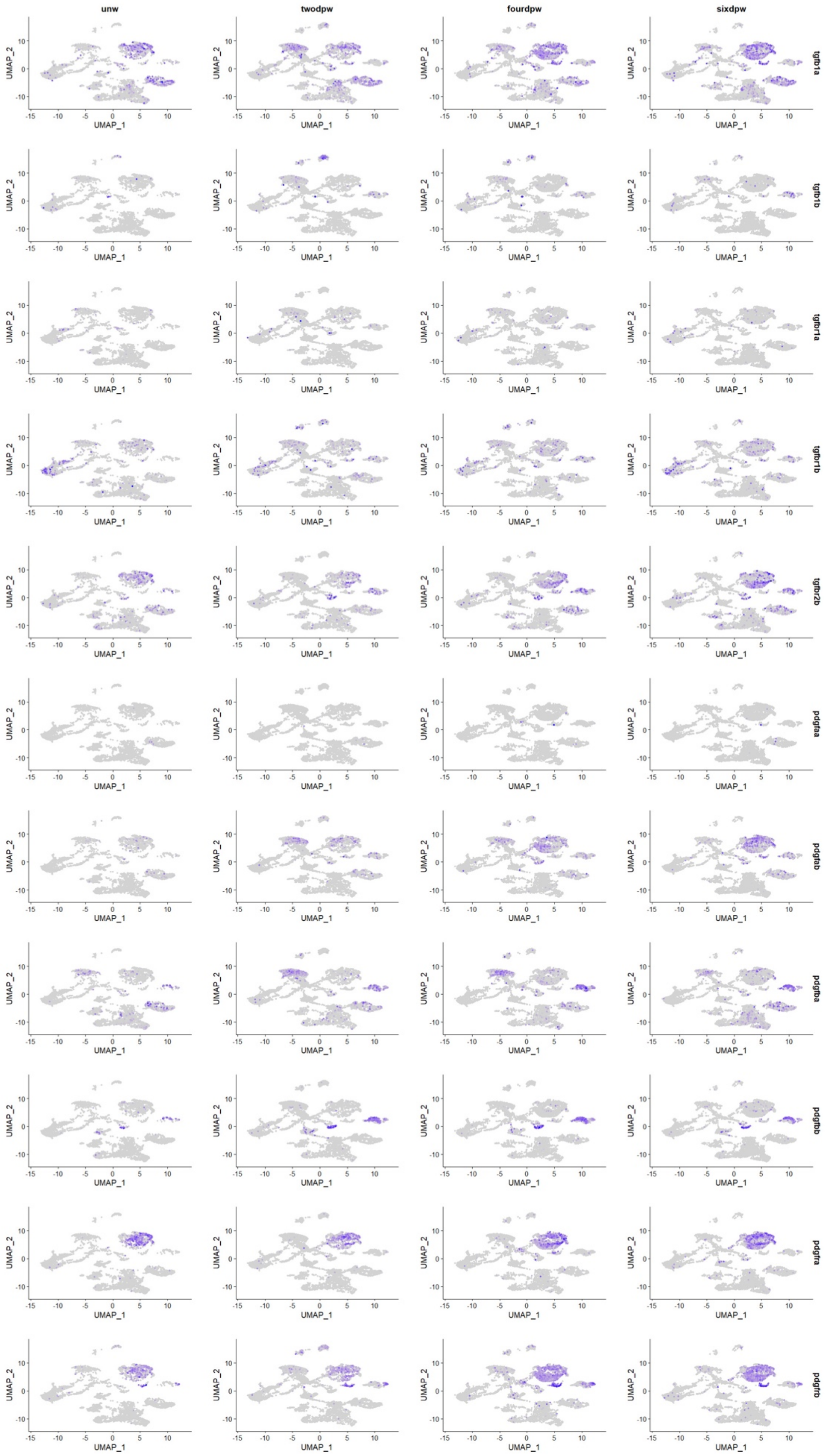
unw
twodpw
fourdpw
sixdpw
tgfb1a
tgfb1b
tgfbr1a
tgfbr1b
tgfbr2b
pdgfaa
pdgfab
pdgfba
pdgfbb
pdgfra
pdgfrb
UMAP_1
UMAP_2

S14 Fig.

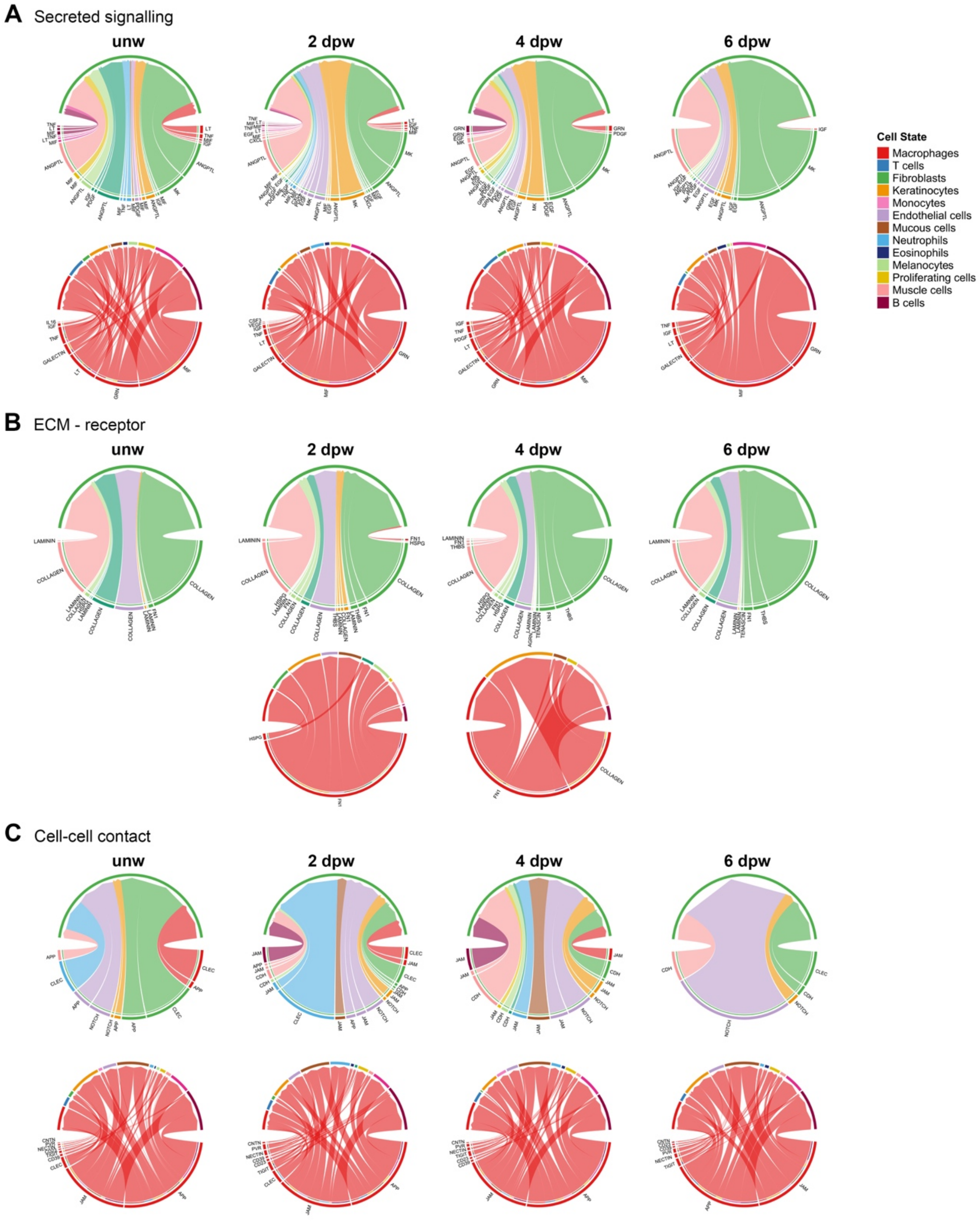
A Secreted signalling
B ECM - receptor
C Cell-cell contact
unw
2 dpw
4 dpw
6 dpw
Cell State
Macrophages
T cells
Fibroblasts
Keratinocytes
Monocytes
Endothelial cells
Mucous cells
Neutrophils
Eosinophils
Melanocytes
Proliferating cells
Muscle cells
B cells

S15 Fig.

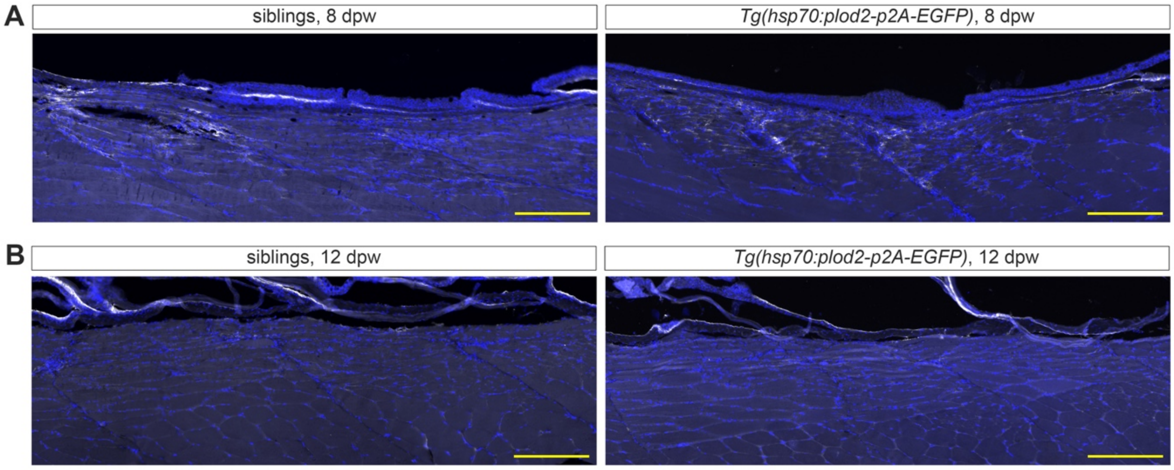
A
siblings, 8 dpw
Tg(hsp70:plod2-p2A-EGFP), 8 dpw
B
siblings, 12 dpw
Tg(hsp70:plod2-p2A-EGFP), 12 dpw

S16 Fig.

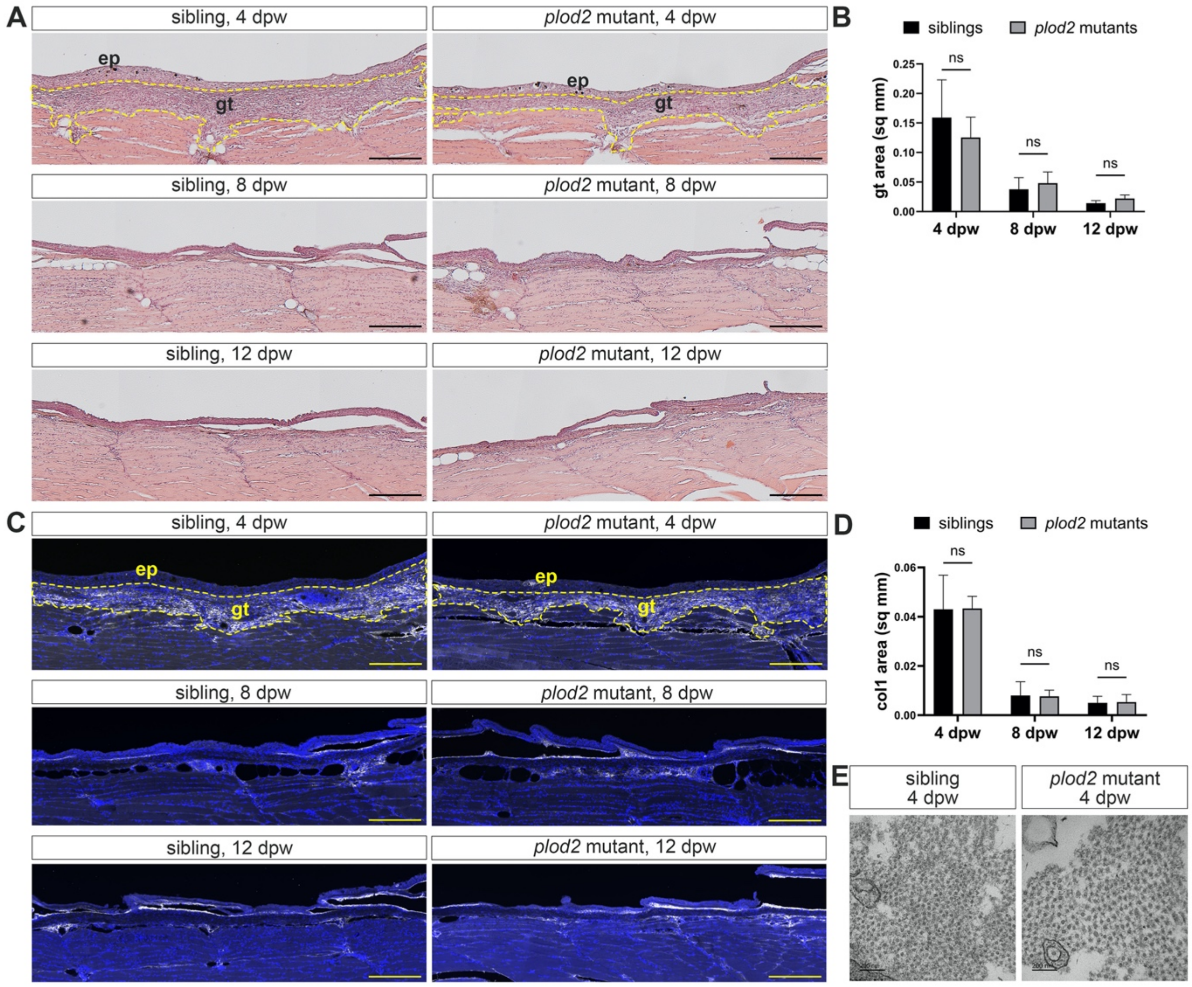

A
sibling, 4 dpw
*plod2* mutant, 4 dpw
ep
gt
sibling, 8 dpw
*plod2* mutant, 8 dpw
sibling, 12 dpw
*plod2* mutant, 12 dpw
B
siblings
*plod2* mutants
gt area (sq mm)
0.25
0.20
0.15
0.10
0.05
0.00
ns
4 dpw
8 dpw
12 dpw
C
sibling, 4 dpw
*plod2* mutant, 4 dpw
ep
gt
sibling, 8 dpw
*plod2* mutant, 8 dpw
sibling, 12 dpw
*plod2* mutant, 12 dpw
D
siblings
*plod2* mutants
col1 area (sq mm)
0.06
0.04
0.02
0.00
ns
4 dpw
8 dpw
12 dpw
E
sibling
4 dpw
*plod2* mutant
4 dpw
